# Aspartate Transaminases Are Dispensable for Pancreatic Development and Pancreatic Cancer Progression

**DOI:** 10.64898/2025.12.17.694916

**Authors:** Julia Ugras, Noah Nelson, Samuel A. Kerk, Damien Sutton, Lin Lin, Peter Sajjakulnukit, Teresa Dombrowski, Gillian Davidson, Brooke Lavoie, Dominik Awad, Alberto Olivei, Wei Yan, Christopher Strayhorn, Megan Radyk, Matthew Perricone, Marina Pasca di Magliano, Filip Bednar, Timothy Frankel, Yatrik M. Shah, Costas A. Lyssiotis

## Abstract

Pancreatic ductal adenocarcinoma (PDA) is the third leading cause of cancer-related deaths in the United States. This is due in part to the limited availability of effective treatment options for patients, highlighting a significant need for new targets and approaches. Deregulated metabolism is a hallmark feature of PDA that has gained attention as a promising inroad for therapy. The aspartate transaminases (<u>g</u>lutamate <u>o</u>xaloacetate transaminases, cytosolic GOT1 and mitochondrial GOT2) have several important metabolic functions, including maintaining energy and redox balance and generating aspartate, an essential building block in protein and nucleotide biosynthesis. Previous studies of GOT proteins in preclinical tumor transplant models have yielded conflicting results regarding the requirement of GOT1 and GOT2 for PDA tumor growth. To assess the role of GOT proteins in tumor development and tumor maintenance, we generated conditional knockout mice for *Got1* and *Got2* and crossed these into pancreas-specific models. Whereas loss of either *Got* does not impact pancreas development, double *Got1* and *Got2* knockout results in markedly reduced pancreas size and cellularity without overtly impacting endocrine or exocrine function. In genetically engineered cancer models, single *Got* loss does not impact lesion formation, tumor size, animal survival, or the composition of the tumor microenvironment. Identical results were also observed in orthotopic allograft mouse models. Together, these findings add to a growing body of work illustrating the adaptability of metabolism in cancer. They also emphasize the importance of model selection, the use of multiple independent models, and the *in vivo* context when studying the role of metabolic programs in cancer.

## Introduction

Pancreatic ductal adenocarcinoma (PDA), the most common form of pancreatic cancer, is projected to be the second leading cause of cancer-related deaths in the United States by 2030^1^. Over half of PDA patients are diagnosed at a late stage of disease when surgery is not possible, and our treatment approaches are largely ineffective. The 5-year survival rate for patients with late-stage disease is only 3%, contrasting the more favorable 5-year survival rate of 44% for those diagnosed when the disease is confined to the pancreas^1–3^. Thus, there is a clear and pressing need to both better detect and intercept in the early stages of disease and develop better treatments for advanced disease.

PDA tumors are noted for their complex cellular environment, which includes a dense stromal reaction that leads to deregulated vasculature, nutrient access, and drug infiltration^4–6^. In this abnormal tumor microenvironment (TME), PDA cells engage in extensive metabolic rewiring to promote their fitness and survival^7^. Many of these mechanisms are driven by oncogenic KRAS^8–11^, which is the signature transforming oncogene in PDA^12–14^. In earlier studies, we found that mutant Kras drives altered glutamine metabolism to support redox balance PDA growth^15^. This phenotype is mediated in part through the rewiring of central carbon metabolism via the glutamate-oxaloacetate transaminases (GOT), cytosolic GOT1 and mitochondrial GOT2.

The GOTs serve many functions in cellular metabolism (**Figure 1A**). They catalyze the reversible, substrate-driven transamination of glutamate and oxaloacetate into aspartate and α-ketoglutarate. Classically, the GOTs are most well known for their role in the malate-aspartate shuttle (MAS), an evolutionarily conserved metabolic pathway that maintains redox and energy balance between the cytosol and the mitochondria. The MAS links cytosolic and mitochondrial energy production through a series of redox reactions that allow NADH to enter mitochondria and support mitochondrial ATP production. In its anabolic role, GOT2 generates aspartate, driven both by facilitated release from the mitochondria and net consumption. GOT1/2 are the only cellular source of aspartate, which is essential for the production of nucleotides and asparagine^15–18^. GOT1 and GOT2 also generate α-ketoglutarate, which can support mitochondrial TCA cycle metabolism and the epigenetic regulation of gene expression.

**Figure 1:**
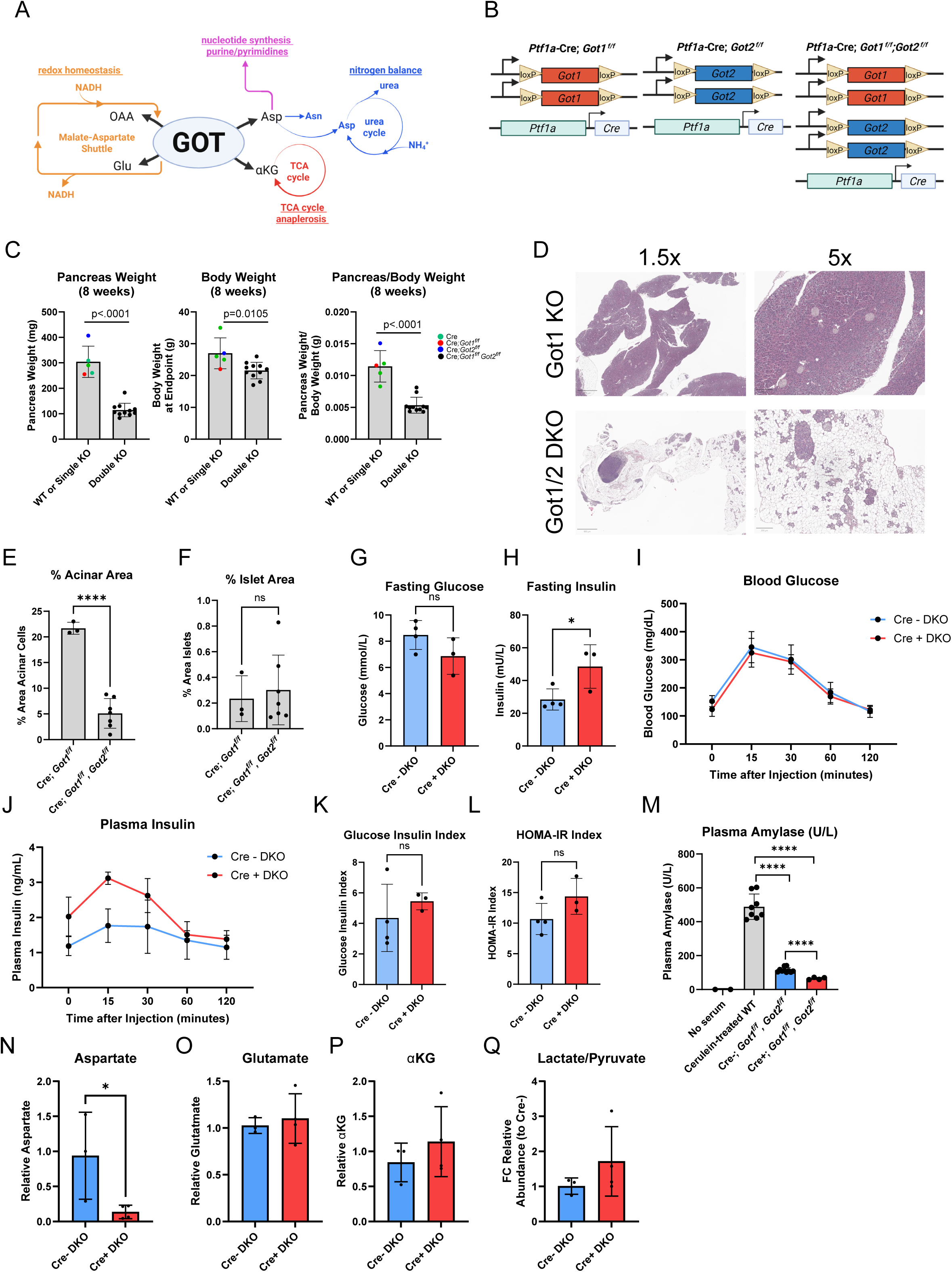
Mouse pancreatic functions are maintained without Got1 or Got2. **A,** Scheme of the glutamate-oxaloacetate transaminases (GOTs) and their roles in cellular metabolism. OAA=oxaloacetate, Asp=aspartate, Glu=glutamine, αKG=α-ketoglutarate, Asn=asparagine. **B,** Scheme of alleles used for *Got* knockout pancreata. *Got1* and/or *Got2* deletion driven by epithelial pancreas-specific Cre recombinase (p48-Cre) on an immunocompetent (C57B/6) background. **C,** Pancreas and body weight of *Ptf1a-Cre*;*Got1^f/f^;Got2^f/f^*double knockout (n=11) mice compared to controls. Controls include 8-week old single KO (n=5) or wild type (WT) mice, as indicated by colored circles. **D,** Representative hematoxylin and eosin (H&E) staining of pancreata from 8-week old Got1 KO and Got1/Got2 knockout mice. Scale bar is 800µm for 1.5x and 200µm for 5x. **E,** Percent acinar and **F,** islet area calculated from H&E images of 8-week old *Ptf1a-Cre*;*Got1^f/f^*(n=3) or *Ptf1a-Cre*;*Got1^f/f^;Got2^f/f^* mice (n=7) mice. **G,** Fasting glucose and **H,** insulin after 5 hour fast in *Got1^f/f^;Got2^f/f^* (n=4) or *Ptf1a-Cre*;*Got1^f/f^;Got2^f/f^*(n=3) female mice. **I,** Blood glucose and **J,** plasma insulin measurements after 1.5g/kg glucose bolus with plasma analysis at 0, 15, 30, 60, and 120 minutes for *Got1^f/f^;Got2^f/f^* (n=4) or *Ptf1a-Cre*;*Got1^f/f^;Got2^f/f^*(n=3) female mice. **K,** Glucose-insulin index and **L,** HOMA-IR index of *Got1^f/f^;Got2^f/f^* (n=4) or *Ptf1a-Cre*;*Got1^f/f^;Got2^f/f^*(n=3) female mice. **M,** Plasma amylase for no serum controls, cerulein-treated wildtype mice (positive controls, n=4), Cre negative *Got1^f/f^;Got2^f/f^*mice (n=5) or *Ptf1a-Cre*;*Got1^f/f^;Got2^f/f^* double knockout mice (n=2). **N-Q,** Relative metabolite abundance, reported as fold change (FC) relative to control, as determined by LC-MS/MS in pancreata of 8-week old *Got1^f/f^;Got2^f/f^* (n=3) or *Ptf1a-Cre*;*Got1^f/f^;Got2^f/f^*(n=4) mice. **N,** aspartate, **O,** glutamate, **P**, α-ketoglutarate (αKG), and **Q**, lactate/pyruvate..

Given their regulation by KRAS and their central functions in cell growth and proliferation, GOT1 and GOT2 have been extensively studied in PDA. For example, we first described how PDA cells route glutamine carbon through a GOT2-GOT1 pathway to maintain redox homeostasis^15^. In this and a subsequent study, we demonstrated that genetic inhibition of GOT1 in PDA cells reduces proliferation and colony formation *in vitro*, sensitizes cells to ferroptosis, and slows the rate of tumor growth in human subcutaneous xenograft models^15,19^. Work from our group on GOT2 revealed that genetic inhibition or knockout of GOT2 reduces proliferation and colony formation *in vitro*, a consequence of elevated NADH reductive stress. However, GOT2 knockout did not slow the rate of tumor growth in subcutaneous and orthotopic human xenograft models^17^. Mechanistically, it was demonstrated that GOT knockout tumors can scavenge aspartate via macropinocytosis to enable tumor growth^20^.

These studies revealed important distinctions between the dependence on GOT metabolism *in vitro* vs *in vivo*. However, transplant models are unable to provide information about the consequences of GOT inhibition in the context of an evolving tumor in the pancreas. To address these shortcomings, we developed a conditional knockout of *Got2* and crossed this into genetically engineered models of pancreatic tumor initiation. We found that *Got2* pancreas-specific knockout did not protect from precancerous lesion formation^17^. A related study using immune competent transplant models illustrated that knockout of *Got2* marked slowed tumor growth in a T cell-dependent manner^21^. It is important to note that these collective studies utilized diverse cell lines and animal models, which may account for the conflicting findings.

More importantly, together, these studies highlight that the field lacks a clear consensus on the role of GOT dependence in PDA.

In this study, we revisit the role of Got1 and Got2 in pancreatic cancer using a battery of state-of-the-art models. Here, we find that the combined loss of Got1 and Got2 leads to reduced pancreas size without overtly impacting function or mouse health. Using these new genetically engineered mouse models, we then demonstrate that loss of either Got1 or Got2 does not impact tumor initiation, growth, or the immune compartment of the TME.

## Results

### Mouse pancreatic functions are maintained without Got1 or Got2

To determine the dependence of normal pancreas development on the GOTs, we crossed *Got1^f/f^* and *Got2^f/f^* mice with the pancreas-specific *Ptf1a*-*Cre* to generate a conditional knockout of *Got1* and *Got2* (**Figure 1B**). We previously published that the single knockout of *Got2* in the pancreas had no observable impact on pancreatic morphology^17^, and similarly we did not observe differences between the knockouts and controls. Accordingly, wild type and single knockouts are used as controls for the double knockout studies (**Figure 1C**). While mice with conditional knockout of both *Got1* and *Got2* are viable, their body weight trailed that of their age-matched controls (by ∼5g on average) but otherwise appeared healthy. Further, analysis of pancreata from double knockout mice taken down at 8 weeks of age revealed significantly smaller pancreata compared to single and wildtype animals (**Figure 1C**).

Histological analysis of double knockout pancreata revealed a significant morphological defect when both *Got1* and *Got2* were lost, compared to the loss of either enzyme alone (**Figure 1D**)^17^. Notable were the loss of acinar tissue area and its repopulation with fat. Using hematoxylin and eosin stains for 8-week-old *Ptf1a-Cre*;*Got1^f/f^*(n=3) and *Ptf1a-Cre*;*Got1^f/f^;Got2^f/f^* mice (n=6), we quantified the acinar and islet area of pancreatic samples. We found that the area of acinar cells was significantly reduced in double knockout mice (**Figure 1D-E**) without an impact on the area of islets (**Figure 1F**). This phenotype is also seen when comparing *Ptf1a-Cre*;*Got1^f/f^;Got2^f/f^* and *Got1^f/f^;Got2^f/f^* (Cre negative) pancreata (**Supplementary Figure 1A**). Next, we performed targeted mass spectrometry-based metabolomics on these pancreata. Consistent with the tissue level disturbances, the metabolome was widely changed (**Supplementary Figure 1C**), which may simply be a result of changes to cellular composition of the tissue. In addition, we observed metabolic shifts coherent with GOT knockout, including a decrease in aspartate and indications of NADH reductive stress (**Figure 1N-Q, Supplementary Figure 1D**). Together, these data illustrate that the loss of *Got1* and *Got2* have a significant impact on pancreatic development, and we questioned whether these mice also had defects in pancreatic exocrine and endocrine function.

Using a separate cohort of double knockout or wildtype female mice, we found that after a 5-hour fast, fasted blood glucose was not significantly altered in double knockout or wildtype animals (n=4 for Cre- control; n=3 for double knockout) (**Figure 1G**). However, fasting insulin was slightly elevated in double knockout mice (**Figure 1H**). After fasting, mice were subjected to a glucose tolerance test. Here, mice were provided with 1.5g/kg of glucose via oral gavage, followed by tail vein blood extraction at 0, 15, 30, 60, and 120 minutes. Here, we find that double knockout animals do not have improved or worsened glucose tolerance or insulin sensitivity compared to control mice (**Figure 1I, J**). Further, *Got1* and *Got2* loss show no significant impact on Glucose Insulin Index or the Homeostasis Model Assessment of Insulin Resistance (HOMA-IR) Index (**Figure 1K, L**). These findings indicate that the available endocrine cells in double knockout pancreata are sufficient to retain normal pancreatic endocrine function.

When acinar cells are damaged, amylase is released into the bloodstream^22,23^. Therefore, to evaluate exocrine function, we measured circulating amylase in the blood in a cohort of double knockout mice and Cre-negative controls. We also included cerulein-treated wildtype mice, which develop pancreatic injury and lesion formation, as a positive control. As expected, cerulein-treated mice (n=4) have high amylase in plasma, ∼4-fold above untreated mice of both genotypes (**Figure 1M**). Double knockout mice (n=2) have significantly lower plasma amylase levels than controls (n=5), potentially reflective of the lower acinar content (**Figure 1M**). As a further measure, feces from the double knockout animals are grossly indistinguishable from controls, further supporting that digestion is overtly normal. Together, these data demonstrate the loss of *Got1* and *Got2* from the pancreas causes a severe disturbance in pancreatic morphology, yet these mice are viable and retain normal pancreatic exocrine and endocrine function at the ages examined.

### Knockout of pancreatic Got1 or Got2 does not impact precancerous lesion formation

Due to the prominent tissue-level defects of *Got1* and *Got2* dual knockout, we elected to use a single knockout of *Got1* or *Got2* for subsequent genetic models of pancreatic cancer. To evaluate whether GOT1 or GOT2 affects the development of precancerous lesions, a first step toward PDA, we crossed the *Got* floxed alleles with the “KC” mouse model. This model expresses a mutant Kras allele (LSL*-Kras^G12D/+^*) driven by the pancreas-specific *Ptf1a*-*Cre* to drive the development of pancreatic lesions and early neoplasia that precedes PDA (**Figure 2A**). KC (n=4), KC;*Got1^f/f^* (n=3), KC;*Got2^f/+^* (n=10), and KC;*Got2^f/f^* (n=5) mice were aged to 6 months, and KC (n=4), KC;*Got1 ^f/+^* KC (n=3);*Got1^f/f^* (n=8), KC;*Got2^f/+^* (n=3), and KC;*Got2^f/f^* (n=4) mice were aged to 12 months, which are well established timepoints to assess the presence of pancreatic transformation. At these endpoints, animals were sacrificed, and pancreata were harvested and processed for histological examination. At both the 6-month and 12-month timepoints, there was no significant difference in animal body weight or pancreas weight for either genotype of KC;*Got* mice (**Figure 2B-E**). Histology from these mice revealed similar levels of metaplasia, lesions, and fibrosis across each group (**Figure 2G**). By 12 months of age, some lesions in KC mice can advance to PDA. A cohort of the 12-month pancreata was further analyzed histologically to grade lesions and calculate the percent tumor area in mice where lesions progressed to PDA (KC n=3, KC;*Got1^f/f^* n=5, KC;*Got2^f/f^* n=5). In this cohort, no wildtype KC mice developed tumors by 12 months, and 2 of 5 *Got1* knockout and 1 of 5 *Got2* knockout pancreata evaluated had small areas of PDA. However, grading these 12-month-old KC mice revealed was no significant difference among the groups for any of the lesion grades evaluated (**Figure 2F**). Additionally, we did not observe any obvious tissue-level morphological differences among genotypes (**Figure 2G-H, Supplementary Figure 2C**). Together, these data suggest that loss of *Got1* or *Got2* is not protective against pancreatic precancerous lesion formation.

**Figure 2:**
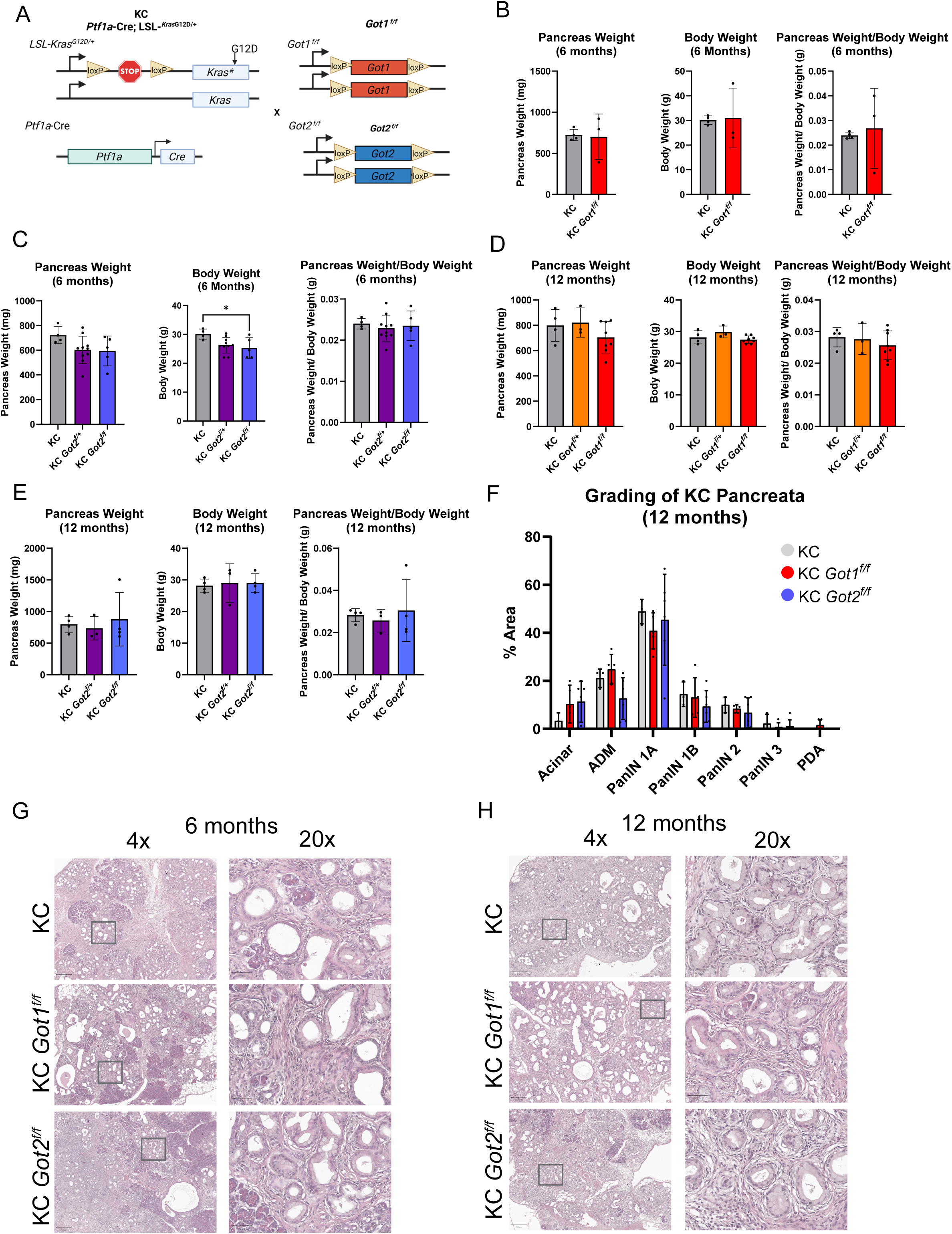
Knockout of pancreatic Got1 or Got2 does not impact precancerous lesion formation. **A,** Scheme of alleles used to study Got in pancreatic precancerous lesions. Got1 or Got2 deletion with expression of mutant Kras (LSL-*Kras^G12D^*) driven by epithelial pancreas-specific Cre recombinase (*p48*-Cre) on an immunocompetent (C57B/6) background (KC-Got1 and KC-Got2). **B,** Pancreas and body weight of KC (n=4) and KC;*Got1^f/f^* (n=3) mice aged to 6 months. **C,** Pancreas and body weight of KC (n=4), KC;*Got2^f/+^* (n=10), and KC;*Got2^f/f^* (n=5) mice aged to 6 months. **D,** Pancreas and body weight of KC (n=4), KC;*Got1 ^f/+^* KC (n=3);*Got1^f/f^* (n=8) mice aged to 12 months. **E,** Pancreas and body weight of KC (n=4), KC;*Got2^f/+^* (n=3), and KC;*Got2^f/f^* (n=4) mice aged to 12 months. **F,** Histological grading of pancreata of KC (n=3), KC;*Got1^f/f^* (n=5), and KC;*Got2^f/f^*(n=5) mice aged to 12 months. **G,** Representative hematoxylin and eosin (H&E) staining of pancreata from 6- and **H,** 12-month aged KC mice. Scale bar is 250µm for 4x and 50µm for 20x images.

### Loss of Got1 or Got2 does not improve survival in genetically engineered models of pancreatic cancer

Next, we assessed the role of the GOT enzymes in PDA using the most well established genetically engineered mouse model of PDA progression. In the KPC model, mice are engineered to express a mutant Kras G12D allele and a mutant TP53 allele (*Tp53 ^R17H2/+^)* driven by the pancreas-specific *Ptf1a-Cre* (**Figure 3A**). This drives pancreatic cancer formation that advances to PDA. To this, we crossed our *Got1* or *Got2* floxed mice. Cohorts of at least 5 mice from each genotype were followed until they died naturally or required sacrifice due to being moribund: KPC (n=6), KPC;*Got1^f/+^* het (n=11), KPC;*Got1^f/f^* null (n=5), KPC;*Got2^f/+^* het (n=12), or KPC;*Got2^f/f^* null (n=11). Among this large cohort of animals, we did not observe significant survival differences among any of the genotypes (**Figure 3B-C**). The mean mortality for each genotype was within a range of 17 days: KPC 134.3 days, KPC;*Got1^f/+^* 128.2 days, KPC;*Got1^f/f^* 145.8 days, KPC;*Got2^f/+^* 133.5 days, KPC;*Got2^f/f^* 143.7 days. These mice were euthanized at various timepoints based upon onset of moribundity; however, within these timepoints, we did not observe any major aberrations in tumor or body weight among our genotypes (**Supplementary Figure 2A-B**). Additionally, we did not observe differences in tumor morphology among genotypes (**Figure 3D**). Together, these data indicate that loss of *Got1* or *Got2* does not improve survival in this genetically engineered mouse model of PDA.

**Figure 3:**
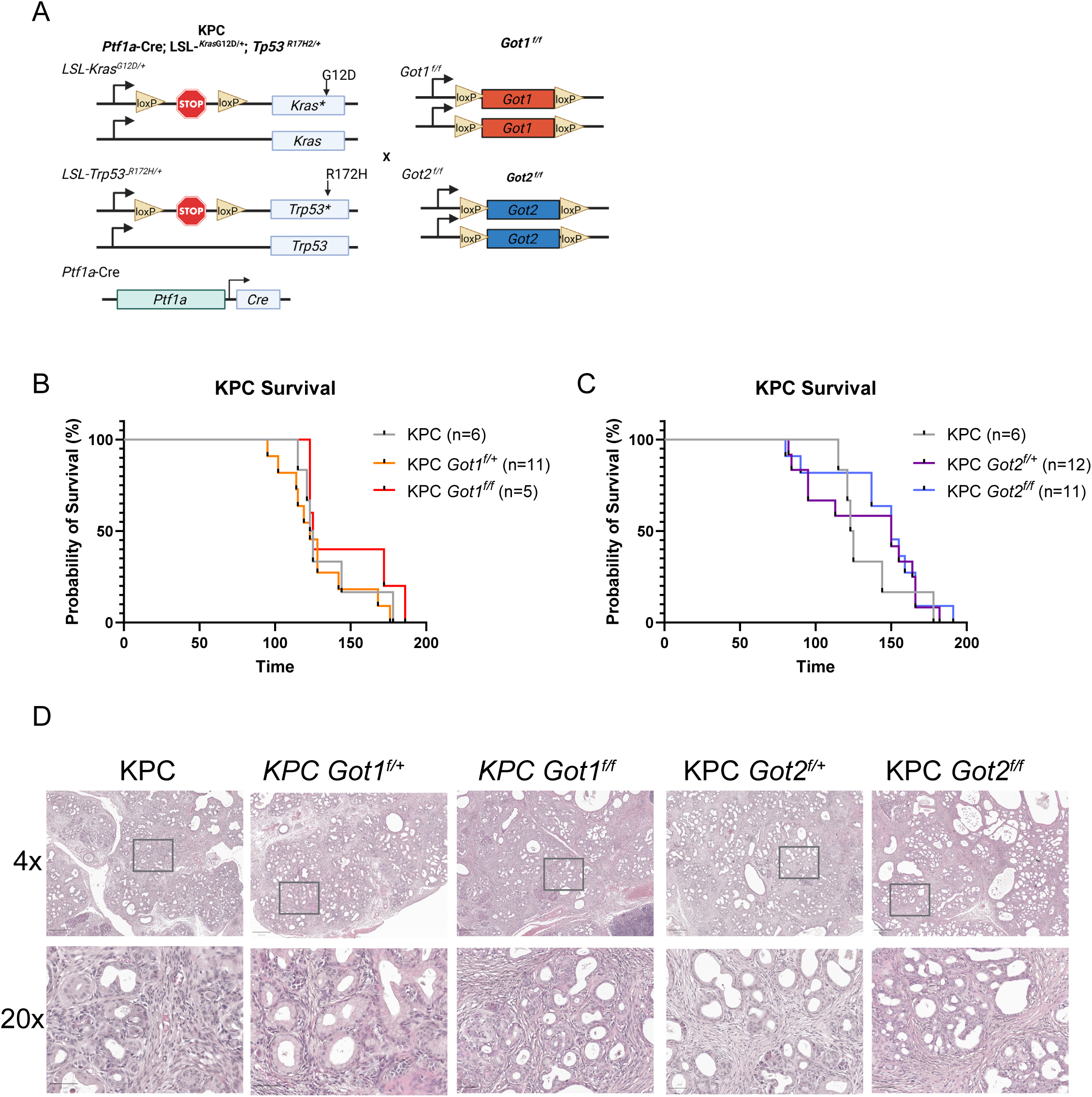
Loss of Got1 or Got2 does not improve survival in genetically engineered models of pancreatic cancer. **A,** Scheme of alleles used to study Got in pancreatic cancer. Got1 or Got2 deletion with expression of mutant Kras (LSL-*Kras^G12D^*) and mutant p53 (LSL-*Tp53 ^R17H2/+^)* driven by epithelial pancreas-specific Cre recombinase (*p48*-Cre) on an immunocompetent (C57B/6) background (KPC-Got1 and KPC-Got2). **B,** Kaplan-Meier survival curves for overall survival of KPC (n=6), KPC;*Got1^f/+^* (n=11), and KPC;*Got1^f/f^* (n=5) mice. **C,** Kaplan-Meier survival curves for overall survival of KPC (n=6), KPC;*Got2^f/+^* (n=12), and KPC;*Got2^f/f^* (n=11) mice. **D,** Representative hematoxylin and eosin (H&E) staining of pancreata from KPC mice. Scale bar is 250µm for 4x and 50µm for 20x images.

### Got1 or Got2 loss in the endogenous KPC tumor model does not alter the composition of immune and epithelial compartments in the tumor microenvironment

A recent study illustrated that *Got2* knockout in syngeneic, orthotopic PDA models reduces tumor size and that this is dependent on the activity of T cells^21^. Although we did not observe an effect on tumor size from either *Got1* or *Got2* loss in our KPC models, we next sought to interrogate whether T cell quantity or subtype was affected by the knockout of *Got1* or *Got2*. Here, we conducted multiplex immunofluorescence (IF) staining on KPC, KPC;*Got1^f/+^*, KPC;*Got1^f/f^*, KPC;*Got2^f/+^,* and KPC;*Got2^f/f^* tumors (n=5 per group). Our multiplex IF panel focused on the quantification of T cells, macrophages, and epithelial cells in these KPC tumors. This included staining for DAPI (nuclei), F480 (macrophages), CD3 (T cells), CD8 (cytotoxic T cells), PDL1 (immune evasiveness), CK19 (epithelial cells), and FoxP3 (Tregs) (**Figure 4A, Supplementary Figure 2C**). These data were then quantified using the inForm software^24–27^. This enabled us to calculate cellular phenotypes and quantities. In the KPC model without disruption of *Got1* or *Got2*, most of the evaluated cell types were epithelial cells and macrophages, with few T cells; data that align with the previous analyses of KPC tumors ^28,29^(**Figure 4B-U**). Most of the macrophages and epithelial cells were PDL1-, which corresponds with PDA tumors being largely unresponsive to single-agent treatment with PDL1 immunotherapies in the clinic (**Figure 4F, I, P, S**)^30^. Compared to macrophages, T cells were in very low abundance (∼2% of all cells), again in line with human PDA and previous analyses of KPC tumors (**Figure 4E, O**)^31^. Compared to the *Got1* and *Got2* knockout tumors, there were no meaningful differences in epithelial or immune cell populations across tumors. This includes no statistical difference amongst the quantity of different T cell populations, including CD3+/CD8-/FoxP3-helper T cells, and CD3+/CD8+/FoxP3-effector CD8+ T cells (**Figure 4B-E, L-O**). We did, however, detect a minor increase in CD3+/CD8-/Fox3P+ Tregs in KPC;*Got1^f.f^* and KPC;*Got2^f/+^* mice, however, these Tregs make up <0.4% of cells (**Figure 4C, M**). We also did not observe a difference in F480+/PDL1+ or F480+/PDL1-macrophages or CK19+/PDL1+ or CK19+/PDL1-epithelial cells (**Figure 4F-K, P-U**). Together, these data demonstrate that *Got1* or *Got2* loss has minimal impact on the immune and epithelial cellular compartments of the TME.

**Figure 4:**
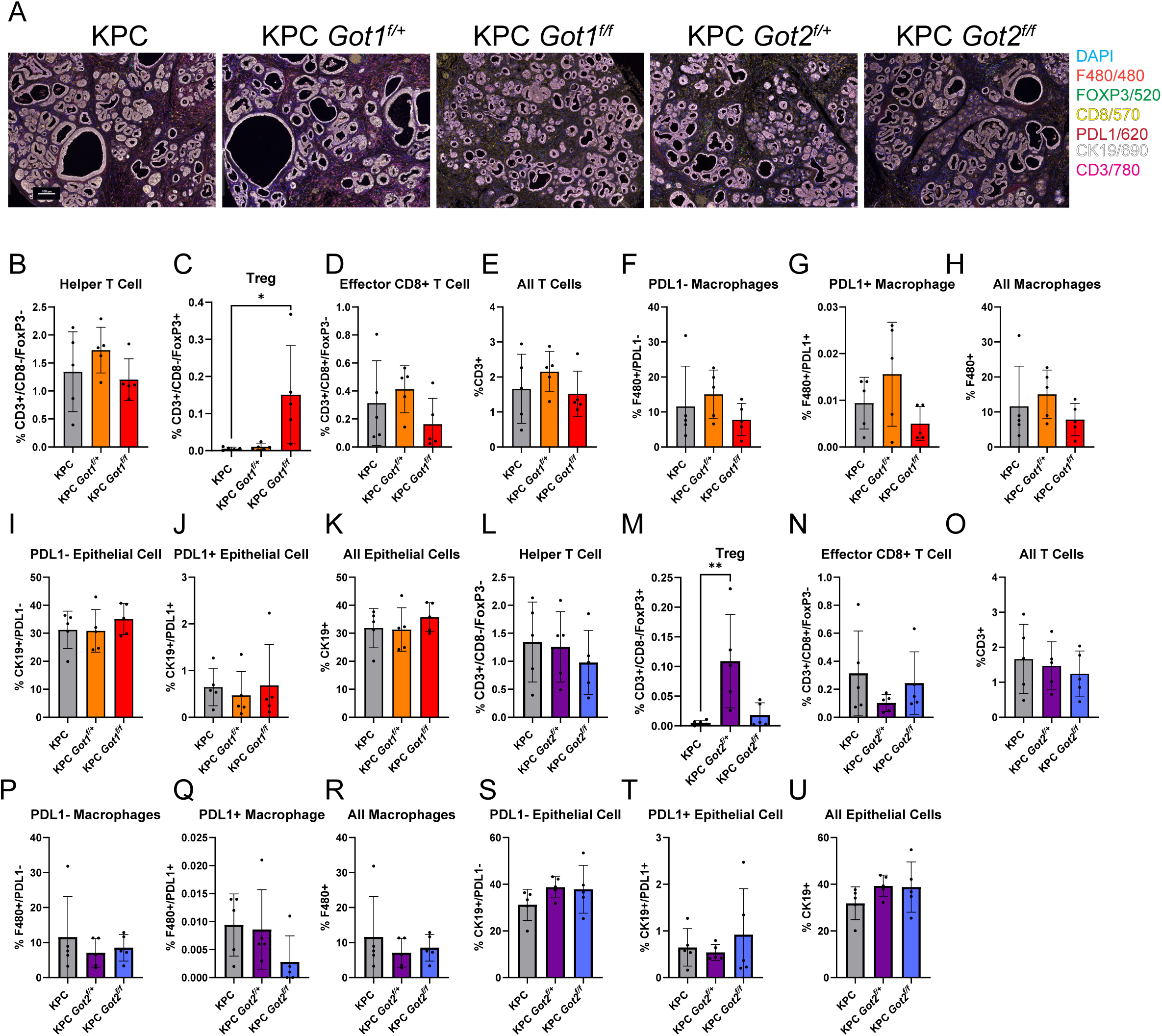
Got1 or Got2 loss does not alter the composition of immune and epithelial compartments in the tumor microenvironment. **A,** Representative multiplex IF images of KPC, KPC;*Got1^f/+^*, KPC;*Got1^f/f^*, KPC;*Got2^f/+^,* and KPC;*Got2^f/f^* mice. Scale bar is 100µm. **B, C, D, E,** T cell populations identified by multiplex IF of KPC, KPC;*Got1^f/+^*, and KPC;*Got1^f/f^* mice (n=5 per group). **F, G, H,** Macrophage populations identified by multiplex IF of KPC, KPC;*Got1^f/+^*, and KPC;*Got1^f/f^* mice (n=5 per group). **I, J, K,** Epithelial cell populations identified by multiplex IF of KPC, KPC;*Got1^f/+^*, and KPC;*Got1^f/f^*mice (n=5 per group). **L, M, N, O,** T cell populations identified by multiplex IF of KPC, KPC;*Got2^f/+^,* and KPC;*Got2^f/f^* mice (n=5 per group). **P, Q, R,** Macrophage populations identified by multiplex IF of KPC, KPC;*Got2^f/+^,* and KPC;*Got2^f/f^* mice (n=5 per group). **S, T, U,** Epithelial cell populations identified by multiplex IF of KPC, KPC;*Got2^f/+^,* and KPC;*Got2^f/f^* mice (n=5 per group).

### PDA cells adapt their metabolism to counter GOT loss

The tissue complexity and limited number of cancer cells in the autochthonous PDA tumors (∼30%, **Figure 4U**) presents a challenge to studying metabolic reprogramming. Thus, we utilized syngeneic KPC cell line-based models to study the cell-autonomous impacts of GOT1 and GOT2. We used CRISPR/Cas9 to knock out *Got1* and *Got2* in the MT3 and 7940B KPC cell lines (**Figure 5A**). Next, in standard cell culture media, we evaluated whether the loss of either enzyme alone affects proliferation of pooled populations, relative to cells harboring a non-targeting sgRNA (sgNT). *Got* knockout did not have a negative effect on proliferation, with a trend toward faster proliferation in the Got1 sg1 condition in both cell lines (**Figure 5B,C**).

**Figure 5:**
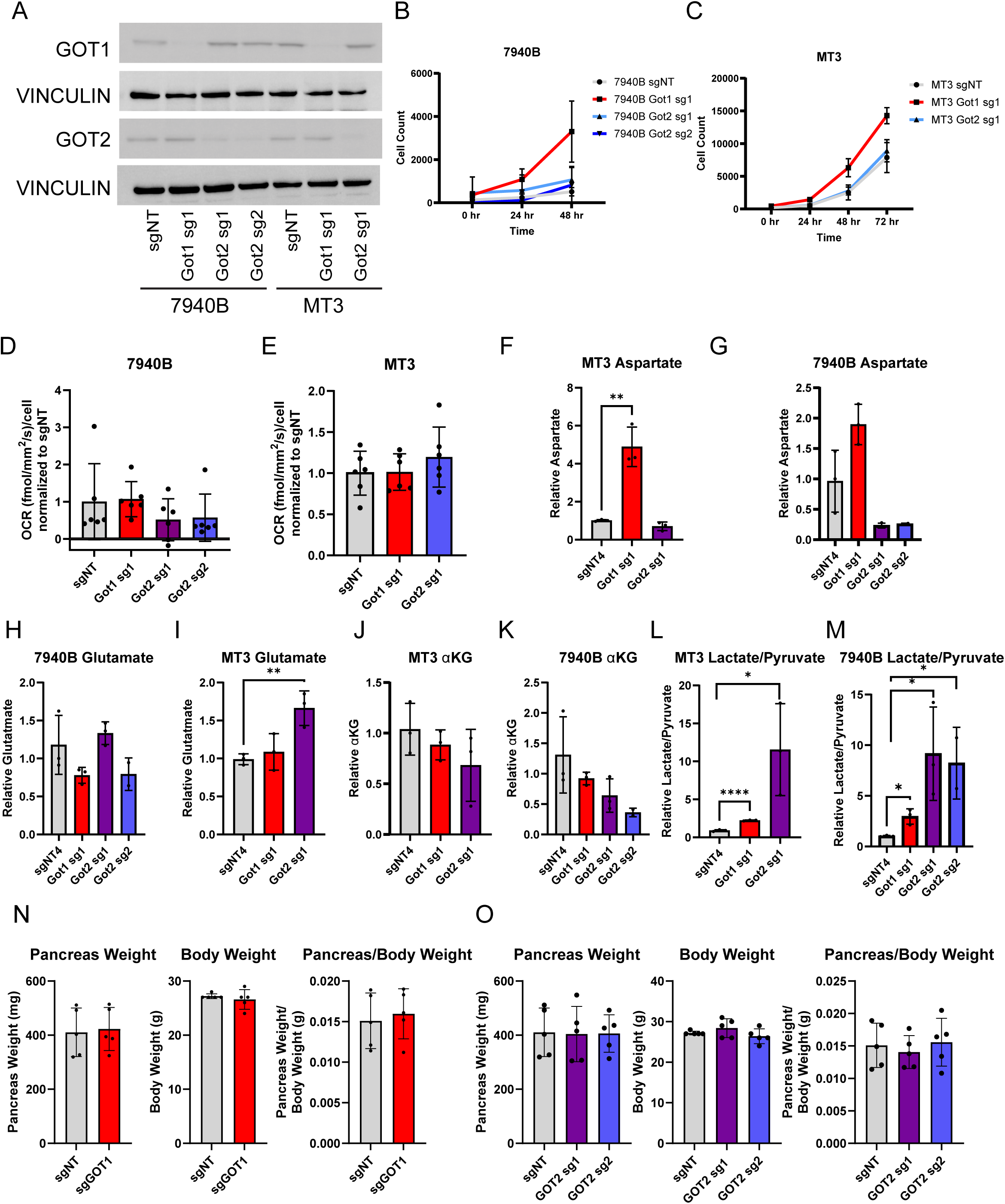
PDA cells adapt their metabolism to counter GOT loss. **A,** Western blot analysis of murine KPC pancreatic cancer cell lines (7940B and MT3) stably expressing a non-targeting sgRNA (sgNT) or with Got1 or Got2 KO. Two independent sgRNAs against GOT2 are designated sg1 and sg2. **B,** Cell count of 7940B and MT3 cell lines with Got1 or Got2 KO over 48-72 hours. Each clone plated with 5-10 replicates. **C,D,** Oxygen consumption rate (fmol/mm^2^/s) of 7940B and MT3 cell lines with Got1 or Got2 KO normalized to cell number. Each clone plated with 6 replicates. **F-M**, Relative intracellular metabolite abundance, reported as fold change (FC), as determined by LC-MS/MS in MT3 and 7940B murine KPC cells. **F,G** aspartate, **H,I,** glutamate, **J,K**, α-KG (α-ketoglutarate), **L,M**, lactate/pyruvate. **N,O,** Pancreas and body weight of mice at endpoint after orthotopic implantation of pancreatic cancer cells (n=5 per group).

Next, we assessed metabolic programs in these cells by measuring oxygen consumption rates and changes to the metabolome by mass spectrometry-based metabolomics. While loss of *Got1* or *Got2* did not negatively oxygen consumption rate (**Figure 5D,E**), we observed widespread changes to both the intracellular and extracellular metabolomes (**Supplementary Figure 3-6**). Notably, aspartate levels tracked with the canonical directionality of the GOTs, with aspartate levels increasing in GOT1 knockouts and aspartate levels decreasing in the GOT2 knockouts (**Figure 5F,G**). Changes to levels of the other GOT substrates were not consistent (**Figure 5H-K**), in line with our previous observations^15,17,19^, and the buffering of glutamate and α-ketoglutarate levels by many other enzymes. We also observed indicators of NADH reductive stress, including a significant increase in the lactate to pyruvate ratio (**Figure 5L,M**).

We then wanted to determine if these metabolic changes would impact tumor growth using an orthotopic, immunocompetent model of PDA. This also allowed us to compare if GOT knockout impacted the take or growth of established tumors, relative to our observations on tumor development and progression using the autochthonous models. To this end, we implanted our Got knockout 7940B KPC cells into the pancreas of B6 mice. Analysis of tumor weight at endpoint revealed that neither *Got1* nor *Got2* loss had a deleterious impact on tumor growth (**Figure 5N,O**). The results from this experiment provide data from a third distinct, immune competent model of PDA, which all collectively illustrate that the GOTs are dispensable in pancreatic tumorigenesis, tumor growth, and tumor progression. They also serve to further underscore the adaptability of pancreatic tumor metabolism.

## Discussion

The concentration of aspartate in the serum is lower than that of all other amino acids, typically low to sub-micromolar. Rather, as a non-essential amino acid, aspartate is created in cells by the GOTs as needed. This suggests that excretory, protein-producing cells should require GOT activity. Single loss of either *Got1* or *Got2* in the pancreas, using the pancreas-specific Ptf1a-Cre, has no apparent impact on pancreatic formation or function. Canonically, mitochondrial GOT2 makes aspartate, which is released into the cytosol by the mitochondrial aspartate-glutamate transporter ARALAR. In the cytosol, GOT1 can act on aspartate and α-ketoglutarate to make glutamate and oxaloacetate. In the absence of GOT2, we speculate that GOT1 runs the canonically reverse reaction to produce aspartate. Indeed, when we fully block aspartate production with dual, pancreas-specific loss of *Got1* and *Got2*, the number of digestive enzyme-producing acinar cells in the pancreas is severely restricted, leading to a much smaller pancreas. Despite this, mice with Got-less acinar cells are viable and do not exhibit overt challenges to digestion, with the potential exception of being slightly smaller in weight. This suggests that a mere fraction of normal pancreas is sufficient to retain normal pancreas function, but more work is necessary to delineate the specific needs of a functioning pancreas.

We speculate that the deleterious impact of Got loss on the exocrine pancreas results from a differentiation defect limiting acinar cell maturation or via cell death or mature acinar cells limited in aspartate. In contrast to acinar cells, total islet area was unaffected by loss of the *Got*s, and the loss of these enzymes did not significantly impact glucose or insulin sensitivity. However, Ptf1a-Cre is less penetrant in the endocrine lineage, and some *Got* expression remains. As such, our observations instead reveal that the major structural impacts to the exocrine pancreas do not appear to impact endocrine function.

Beyond its role in protein biosynthesis, aspartate is also an essential building block for nucleotides and other amino acids, highlighting the potential importance of the GOTs for the biosynthetic needs of proliferating cells^16,18^. Further, the action of the GOTs within the MAS are also important mediators of redox balance through their downstream impacts on NADH/NAD^+^ ratios^15^. Together, the physiological roles of the GOTs and the MAS suggest a potential importance of these pathways in cancer cells, where redox imbalances and biosynthetic needs are paramount^7,15,32^. Despite these important functions, we find that the pancreas-specific loss of *Got1* or *Got2* does not protect against precancerous lesions or reduce PDA tumor size using orthotopic and genetic models of pancreatic cancer *in vivo*.

Our data herein add to a growing body of work on the role of GOTs in PDA and include some conflicting observations. Over a decade ago, we demonstrated that PDA cells rely on the metabolism of glutamine-derived carbon through the activity of GOT1 in a pathway regulated by mutant KRAS^15^. Using human cell culture models *in vitro* and immunocompromised xenograft mouse models, we showed that genetic GOT1 inhibition slows PDA growth and leads to oxidative stress, which sensitizes cells to ferroptosis^19^.

We also explored the role of GOT2 in pancreatic cancer. Using human cell line models *in vitro*, we found that genetic inhibition of GOT2 reduces PDA cell proliferation^17^. In contrast, after implanting human PDA cell lines subcutaneously or orthotopically into nude mice, we found that genetic inhibition of *GOT2* does not impair tumor growth. In the genetically engineered KC model, *Got2^f/f^*knockout had no impact on disease progression. Subsequent studies *in vitro* using human PDA cell lines revealed that GOT2 knockdown led to NADH reductive stress, which inhibited proliferation. Reversal of the NADH burden restored proliferation. Indeed, we demonstrated conditioned media from cancer associated fibroblasts could rescue GOT2 proliferative defects in vitro, and this activity was attributed to pyruvate, potentially explaining why *Got2* loss does not affect tumor growth^17^. A related study also found that GOT2 knockout had no impact on PDA tumor growth using human cell lines in immune compromised animals^20^. Further, they demonstrated that aspartate auxotrophic tumors (double GOT1-GOT2 knockout), while slower growing, maintained aspartate levels through macropinocytosis, providing important new mechanistic insights into how tumors cope with GOT loss.

In contrast to the GOT2 results using human cell lines in immune compromised mice described above, it was reported that *Got2* knockout using murine KPC cells and syngeneic, orthotopic transplantation resulted in markedly reduced tumor growth^15,19^. Tumor growth inhibition was ascribed to tumor-infiltrating T cells and a decrease in immunosuppressive macrophages in this immune competent model.

Building on our previous work studying GOTs in human cell lines and immune compromised transplant models, here we utilized mouse models pancreatic cancer development and progression *in vivo*, along with orthotopic animal models and cell line models. We find that murine tumor development, progression, and established tumor models appear largely, if not completely, unaffected by *Got1* or *Got2* loss. These results are harmonious with previous work on GOT2 in human models^17^ ^20^ yet contrast those from another mouse study^21^. These new results also stand in contrast to our previous work on GOT1 in human models^15,19^. It is likely that we can fully reconcile these discrepancies by simply considering the features of the model systems employed. Such results also serve to highlight concerns with reproducibility amongst scientific studies, which may not in fact be issues with reproducibility but rather a consequence of the myriad variables associated with studying cancer in animals.

Finally, the observation that PDA tumors can develop, grow, and progress without Got1 or Got2 highlights the adaptability of cancer metabolism. Indeed, a wealth of studies detail the metabolic flexibility of PDA, including their ability to engage various recycling and scavenging pathways, as well as intratumoral metabolic crosstalk pathways, to facilitate growth and survival. For example, our metabolomics data and that of others indicates isoform compensation for aspartate biosynthesis. Additionally, aspartate auxotrophic tumors overcome dual GOT loss by utilizing macropinocytosis to scavenge for environmental aspartate^20^. Single Got knockout will also disable the malate-aspartate shuttle and thus lead to NADH/NAD+ redox imbalance. Our previous work on fibroblast-derived pyruvate release may provide a mechanism to overcome NADH stress^17^. Future work will be necessary to study these metabolic compensatory mechanisms.

## Author Details and Contributions

J.U., N.N., S.A.K., Y.S., and C.A.L. designed the study. J.U. and C.A.L. wrote the manuscript. J.U. and N.N collected data for the bulk of experimental studies. J.U. prepared manuscript figures. S.K., D.S., T.D., G.D., M.P., M.R., D.A., A.O., L.L., P.S., W.Y., and C.S conducted experiments. A.O., P.S., and W.Y. contributed to data analysis. C.A.L., Y.S., F.B., T.F., and M.P.M. provided resources, funding, and conceptual input for experiments and supervised the research. All authors reviewed and approved the manuscript.

## Acknowledgements

We would like to acknowledge the services provided by the Mouse Metabolic Phenotyping Center-Live at Michigan through their grant U2CDK135066. This research was supported by the NCI (C.A.L. R37CA237421, R01CA248160, R01CA244931). J.U. was supported by T32DK094775.

## Methods

### *Got* CRISPR/Cas9 knockout

The lentiviral vector for CRISPR knockout (KO), lentiCRISPR v2, was obtained from Addgene (plasmid #52961)^33^. Plasmid DNA was expanded by transformation of NEB Stable competent cells (NEB, C3040H) and maxi prepped using the EndoFree Plasmid Maxi Kit (Qiagen, 12362). The cloning vector was prepared by *BsmB*I-v2 (NEB, R0739S) digestion and the ends of linearized vector dephosphorylation executed with Shrimp Alkaline Phosphatase (rSAP) (NEB, M0371S) followed by 1% of agarose gel (Invitrogen, 16500500) purification (Qiagen, 28704). Several guide RNAs from the various mouse CRISPR libraries were selected and screened (https://www.addgene.org/pooled-library/#crispr). Previously published cloning procedures were followed^33,34^. The workflow for generating *Got1* and *Got2* knockout cell lines are detailed here. 1) Sense and antisense guide RNA oligos were synthesized (IDT-Integrated DNA Technologies), phosphorylated (NEB, M0201S), and annealed (ABI Thermal Cycler). The prepped vector and annealed guide RNAs were ligated (NEB, M0202S); 2) 293FT cells were used for transfection; 3) 7940B and MT3 murine pancreatic cancer cell lines were transduced with virus and selected with puromycin for 5 days (Sigma, P8833, 3µg/mL). The best KO guide RNA constructs were selected after Western Blotting and qRT-PR and utilized for *in vitro* and *in vivo* studies. Sequences of mouse guide RNAs for mouse Got1 (NM_010324) and Got2 (NM_010325) are shown here.

**Table 1.**
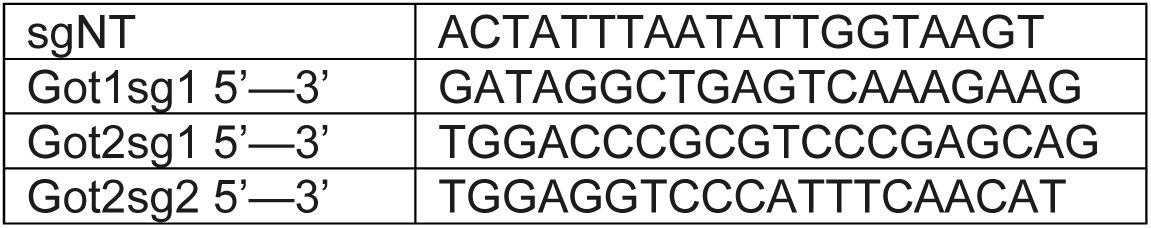
sgRNA sequences.

### Western Blotting

For cells in culture, medium was aspirated, and the wells were washed once with PBS. Thereafter, 50-100 μl of radioimmunoprecipitation assay buffer (Sigma-Aldrich, R0278) to which phosphatase and protease inhibitors were added, was transferred to each well to lyse the cells. Lysis and the collection of the lysates were completed on ice. Following a 5- to 10-minute incubation on ice, lysates were collected into 1.5 ml Eppendorf tubes and centrifuged at 4 °C for 10 min at 18,000*g* to extract the sample supernatant. Protein concentration of the samples for western blot analysis was measured using Pierce BCA Protein Assay Kit (ThermoFisher, 23227) according to the manufacturer’s instructions. For the running step, samples were loaded at 20 μg protein per lane along with the SeeBlue Plus2 protein ladder (Invitrogen, LC5925) and run at 120V on an Invitrogen NuPAGE 4–12% Bis-Tris gel for 1 hour (ThermoFisher, NP0336BOX). Thereafter, the separated proteins were transferred to methanol-activated PVDF membranes (Millipore) at 25 V for 1 h. Following this, membranes were immersed in blocking buffer (5% blotting-grade blocker (Bio-Rad, 1706404) in TBS-T solution: tris-buffered saline (Bio-Rad, 1706435) with 0.1% Tween-20 (Sigma-Aldrich, 9005-64-5) for ∼1 h on a plate rocker at room temperature. Next, membranes were washed 3 times with TBS-T at 10 min per wash, immersed in the indicated primary antibodies, and incubated overnight at 4 °C on a plate rocker. The antibodies used were diluted in blocking buffer at dilutions recommended by the manufacturer. The following day, the primary antibody was removed, and the membrane was washed 3 times with TBS-T and on a plate rocker for 5 min per wash. Immediately after, the membrane was incubated for 1 h with gentle rocking at room temperature in the appropriate secondary antibody diluted 1:10,000 in TBS-T. Lastly, the membrane was washed 3 times in TBS-T at 10 min per wash and incubated in chemiluminescence reagent (Clarity Max Western ECL Substrate, 705062) according to the manufacturer’s instructions. Subsequently, blot images were acquired on a Bio-Rad ChemiDoc Imaging System (Image Lab Touch Software version 2.4.0.03). The following primary antibodies were used in this study at a 1:1,000 dilution: GOT1 Abcam 239487, GOT2 Sigma-HPA018139, and Vinculin CST (E1E9V) XP Rabbit mAb #13901. For secondary antibodies, Cell Signaling Anti-rabbit HRP-linked antibody (7074) and Anti-mouse HRP-linked antibody (7076) were used at a 1:10,000 dilution.

### Cell viability assay

Cells were seeded in replicates of 5 or 10 at 500 cells per well and 1000 cells per well in standard growth medium for 24 hours in a white-bottom 96-well plate. The next day, in dark conditions, Cell Titer Glo Reagent (CAT G9242) was diluted 1:3 in PBS and added in a 1:1 volume to the media in each well. Plates were covered from light, shaken for 2 minutes, then incubated at room temperature for 10 minutes to stabilize the signal. Luminescence was promptly recorded on a plate reader.

### Cell proliferation assay

Cells were seeded in replicates of 5 or 10 at 500 cells per well and 1000 cells per well in standard growth medium for 24 hours in a clear-bottom 96-well plate. Cells were then placed on a Biotek Biospa 8 Automated Incubator for continuous incubation and cell counting over 48-72 hours. Cell counts were taken every 24 hours for each well.

### Oxygen consumption rate

Cells were seeded in replicates of 7 at 500 cells per well in standard growth medium for 24 hours in a Falcon clear-bottom 96-well plate. The next day, a Lucid Scientific Resipher was placed on each plate in a sterile incubator for 72 hours to measure oxygen consumption rate and environmental conditions, including humidity. Data was analyzed from the Lucid Lab online tool. Oxygen consumption rate was normalized to cell count for each line, followed by normalization to the corresponding non-targeting sample.

### Metabolomics Sample Preparation

For *in vitro* extracellular (medium) and intracellular metabolomic profiling, PDA cells were seeded in triplicate in a 6-well plate at 150,000 cells per well in growth medium. The cells were incubated for 48 hours. Thereafter, for extracellular metabolites, 200 µl of medium was collected from each well into a 1.5 ml Eppendorf tube, spun to remove cell debris, and to that 800 µl ice-cold 80% methanol was added. For intracellular metabolites, the remaining medium was aspirated, and 1 ml of ice-cold 80% methanol was added to each well on dry ice for 10 min. Each well was scraped and spun down to clarify samples. Thereafter, cell lysates were collected from each well and transferred into separate 1.5 ml Eppendorf tubes. The samples were then centrifuged at 12,000*g*. For each experimental condition, the volume of supernatant to collect for drying with the SpeedVac Vacuum Concentrator (model: SPD1030) was determined based on the protein concentration of the parallel plate.

For pancreas tissue, samples were flash frozen in liquid nitrogen upon collection. Tissue samples of approximately equal weight (∼0.02 g) were collected per sample per experimental group. The tissues were then put into 2 ml Eppendorf tubes with 1 ml of ice-cold 80% methanol (diluted in 20% H_2_O). Metallic beads were added to each tube, and samples were shaken and homogenized on a Retsch TissueLyser II (129251128) in intervals of 30 s until fully homogenized. Samples were then centrifuged at 12,000*g,* and the supernatant was collected for further processing.

### Metabolomics method and analysis

Targeted metabolomics was performed on an Agilent 1290 Infinity II Binary Bio LC coupled with an Agilent 6495d QqQ mass spectrometer. Two methods were utilized in this study on the same instrumentation. For the first method, an Agilent InfinityLab Poroshell 120 HILIC-Z, 2.1 x 150 mm, 2.7uM (p/n 683775-924) was used. Method parameters are as follows: Solvent A is water + 20mM ammonium acetate (pH 9.3) + 5uM medronic acid. Solvent B is acetonitrile. Wash solvent is 1:1:1 water, acetonitrile, and methanol. The solvent gradient is 10% A – 90% B until 1 minute, 22% A – 78% B until 8 minutes, 40% A – 60% B until 12 minutes, 90% A – 10% B until 15 minutes, hold until 18 minutes, 10% A – 90% B until 19 minutes, hold until 23 minutes. Flow rate was 0.4mL/minutes. Column temperature was set to 15 C. Source parameters were optimized for the Agilent dMRM (dynamic multiple reaction monitoring) library. The dMRM library was acquired from Agilent for their HILIC-Z platform and further optimized to screen for 435 targets in both positive and negative ion modes.

Method 2 utilized an Agilent ZORBAX RRHD Eclipse Plus C18, 100 x 2.1mm, 1.8um (p/n, 959758-902) column. Solvent A was water + 0.1% formic acid. Solvent B is methanol + 0.1% formic acid. Wash solvent is 1:1 water and methanol. The solvent gradient is100% A until 3 minutes, 5% A – 95% B until 5 minutes, hold until 7 minutes, 100% A until 8 minutes, hold until 10 minutes. Flow rate is 0.2mL/minutes. Column temperature is set to 45 C. Source parameters reflect method 1. The dMRM acquisition list includes 12 targets optimized by analytical standards in both positive and negative ion modes.

Data from both methods was preprocessed using Agilent MassHunter Workstation Quantitative Analysis for QQQ Version 12.1.

Raw data was normalized by median-centering and then further fold-change analysis was performed relative to control groups to achieve a relative abundance. Student’s T-Tests were performed with an alpha level of 0.1. Heatmaps were generated in Morpheus (Morpheus, https://software.broadinstitute.org/morpheus).

### Mouse studies

Animal studies were performed at the University of Michigan under the supervision of Unit for Laboratory Animal Medicine and in accordance with the Institutional Animal Care and Use Committee (IACUC). All animals used in this study were housed in a pathogen-free environment, and all procedures involving these animals were performed in accordance with the requirements of the University of Michigan Institutional Animal Care & Use Committee (IACUC). Mice were housed at a maximum of five mice per cage in a pathogen-free animal facility with a 12 h:12 h light: dark cycle, with 30–70% humidity and a temperature of 20–23 °C.

### Genetically engineered mouse models

*Ptf1a-Cre, Ptf1a-Cre;*LSL*-Kras^G12D/+^* and *Ptf1a-Cre;*LSL*-Kras^G12D/+^;Trp53^R172H^*^/+^ mice have been described previously^12^. Floxed *Got1^f/f^* and *Got2^f/f^* mice were generated by Ozgene (*Got1* floxed around exon 3; *Got2* floxed around exon 2)^17,35^. PCR genotyping was done for the *Ptf1a-Cre, Kras^G12D/+^*, *Trp53^R172H^*^/+^, *Got1^f/f,^* and *Got2^f/f^* alleles from DNA isolated from mouse tails using standard methodology, using the primers listed below. Littermate controls were used in all experiments, and the sex ratios for each cohort were balanced.

For the double knockout Cre mice, mice were euthanized at 8 weeks to evaluate the pancreas and body weight. For the double knockout Cre mice used in the glucose tolerance test, mice were euthanized 1 week after test completion. Pancreata and plasma were collected for downstream analyses.

Mice in the KC experiment were euthanized at 6 or 12 months of age. Pancreata were collected for downstream analyses.

For the KPC study, mice were euthanized at humane endpoint according to veterinarian recommendations based on disease symptoms or at 20% body weight loss. Pancreata and plasma were collected for downstream analyses.

### Cerulein treatment

Wildtype mice (2 male, 2 female) were injected with cerulein (Caerulein, Sulfated TS# J64320MCR) hourly for a total of 8 times on day one, followed the next day with 2 more injections and euthanasia an hour later. Cerulein was administered at 75ug/kg via intraperitoneal injection. At euthanasia, feces and pancreata were collected for downstream analysis, and blood was collected via cardiac puncture.

### Orthotopic tumor implantation

Male 8- to 12-week-old C57BL/6J mice were obtained from The Jackson Laboratory (strain 000664) and maintained in the facilities of the Unit for Laboratory Animal Medicine (ULAM) under specific pathogen-free conditions. Before tumor cell injection, wildtype or *Got1*-KO or *Got2*-KO mouse cell lines (derived from KPC 7940b) were collected from culture plates according to standard cell culture procedures. The cells were counted, washed once with PBS, and resuspended in a 1:1 solution of serum-free DMEM and Matrigel (Corning, 354234). For the orthotopic surgical procedure, mice were anaesthetized using inhalation isoflurane. The surgical site was sterilized by swabbing with iodine (Povidine-Iodine Prep Pad, PDI, B40600). This was followed by an incision on the left flank using sterilized instruments. Thereafter, the cell lines were injected into the pancreas, and the incision was sutured. Cell injection was as follows: 50,000 cells in 50 μl final volume for orthotopic implantation. Animals were monitored regularly in the post-operative period, and the orthotopic group was euthanized at post-implantation day 18. At endpoint, pancreata were weighed and preserved for further analyses.

### Tissue processing

Mice were euthanized by CO2 asphyxiation or isoflurane overdose, followed by cervical dislocation. Tissue was quickly harvested and fixed at room temperature with zinc formalin fixative (Z-Fix; Anatech Ltd). Fixed tissues were transferred to 70% ethanol after 24 hours. Tissues were processed using a Leica ASP300S Tissue Processor (Leica Microsystems, Inc), paraffin embedded and cut into 5 μm sections. One section was stained for H&E for histological analysis.

### Histology imaging and grading

Imaging was performed automatically using a whole-slide scanner (Vectra Polaris, Akoya Biosciences) or manually using an Olympus BX53F microscope. Whole-slide scanning was performed by the by the University of Michigan Tissue & Molecular Pathology Shared Resource. For tissue grading, at least 3, 20x fields were reviewed by a pathologist and graded in a blinded manner.

### Quantification of islet and acinar area

Hematoxylin and eosin-stained slides were digitally scanned and loaded into QuPath. Total tissue area was identified using a Pixel Classifier for detecting Average Channels with a sigma of 2 and threshold of 227. The spleen was manually removed from analyses. Islets and acinar area were manually annotated. Measurements were exported and analyzed to obtain percent acinar area and percent islet area.

### Opal multiplex IHC staining

Blocks were cut into 5-micron slices and placed onto charged slides for processing. Slides were baked at 60℃ for 1 hour. Slides were subjected to deparaffinization and rehydration, then fixed with formalin. The multiplex staining was performed according to established protocols^24^. Using the OPAL 7 manual kit, the slides underwent six rounds of staining. Slides were prepared for each staining round using AR9 or AR6 Akoya Biosciences antigen retrieval buffer, followed by a primary antibody (F480 from Cell Signaling, cat #70076S;CD3 from Abcam, cat #Ab11089;CD8 from Cell Signaling, cat #98941;PDL1 from eBioscience, cat #14-5982-81;CK19 from DSHB, cat #Ab 2133570;FoxP3 from Cell Signaling, cat #12653). This was followed by secondary antibody stain (ImmPRESS HRP Goat Anti-Mouse IgG Polymer Detection Kit and ImmPRESS HRP Horse Anti-Rabbit IgG Polymer Detection Kit). Slides were counterstained using 4’,6-diamidino-2-phenylindole (DAPI), mounted, cover-slipped, and left to dry overnight. Using the PhenoImager HT (Akoya Biosciences), slides were imaged at 20x in all channels: DAPI, Opal 480, Opal 520, Opal 570, Opal 620, Opal 690, and Opal 780. Composite images were created by automatically merging images from each channel and then taken for further analysis.

### OPAL analysis

Images were analyzed using inForm Cell Analysis software (Perkin Elmer). All 90 images were batch analyzed using a subset of 13 randomly chosen tissue core images. A library was made using single-antigen staining for each fluorophore and used for unmixing and validation of the multiplex fluorescent composite staining, ensuring no spectral overlap between fluorophores. Using the inForm training software, DAPI counterstain was used to determine the size and shape of each nucleus, with a splitting sensitivity of 0.5. After cell segmentation, the following cells were phenotyped using the trainable software after select cells were manually assigned based on single staining criteria: T cells (CD3+), cytotoxic T cells (CD8+), macrophages (F480+), Tregs (FoxP3+), epithelial cells (CK19+). Fluorescent intensity scores of PD-L1, CD8, and FoxP3 were determined, which allowed for complex cell phenotyping using R^24^. After the fluorescent intensity score was determined for PD-L1, CD8, and FoxP3, using R programs in combination with the original cell phenotypes produced by inForm (T cell, APC, EC, and other), complex phenotypes were formulated. Final multiplex fluorescent composite images were reviewed with a trained pathologist to confirm the accuracy of staining and phenotyping.

### Glucose tolerance test

After 5 hours fast, animals were given 50% glucose via oral gavage at 1∼2.0g/kg. Blood samples are collected before and after gavage at time 0, 15, 30, 60, and 120 minutes via tail vein bleeding. Blood levels of glucose were measured using a glucometer (Acucheck, Roche), and plasma levels of insulin are determined using the Millipore rat/mouse insulin ELISA kit. Animals were restrained repeatedly for less than a minute each time while blood samples were collected. HOMA-IR was calculated by multiplying fasting glucose by fasting insulin and dividing by 22.5. Glucose insulin index was calculated by multiplying the area under the curve of glucose and the area under the curve of insulin and 10^-6^.

### Amylase

Plasma obtained via tail vein bleed or cardiac puncture was used to quantify amylase in plasma using the Pointe Scientific Amylase Reagent (Fisher 23-666-111). Following the manufacturer’s instructions, 10 μL of plasma was added to 400uL of reagent, followed by reading on a plate reader at 409nm with 3 readings taken over 2 minutes. To calculate the concentration of amylase in a sample in U/L, the average absorbance per minute was multiplied by the assay volume, then by 1000. This product was divided by the millimolar absorptivity of 2-chloro-p-nitrophenol times the sample volume times the light path.

**Table 2.**
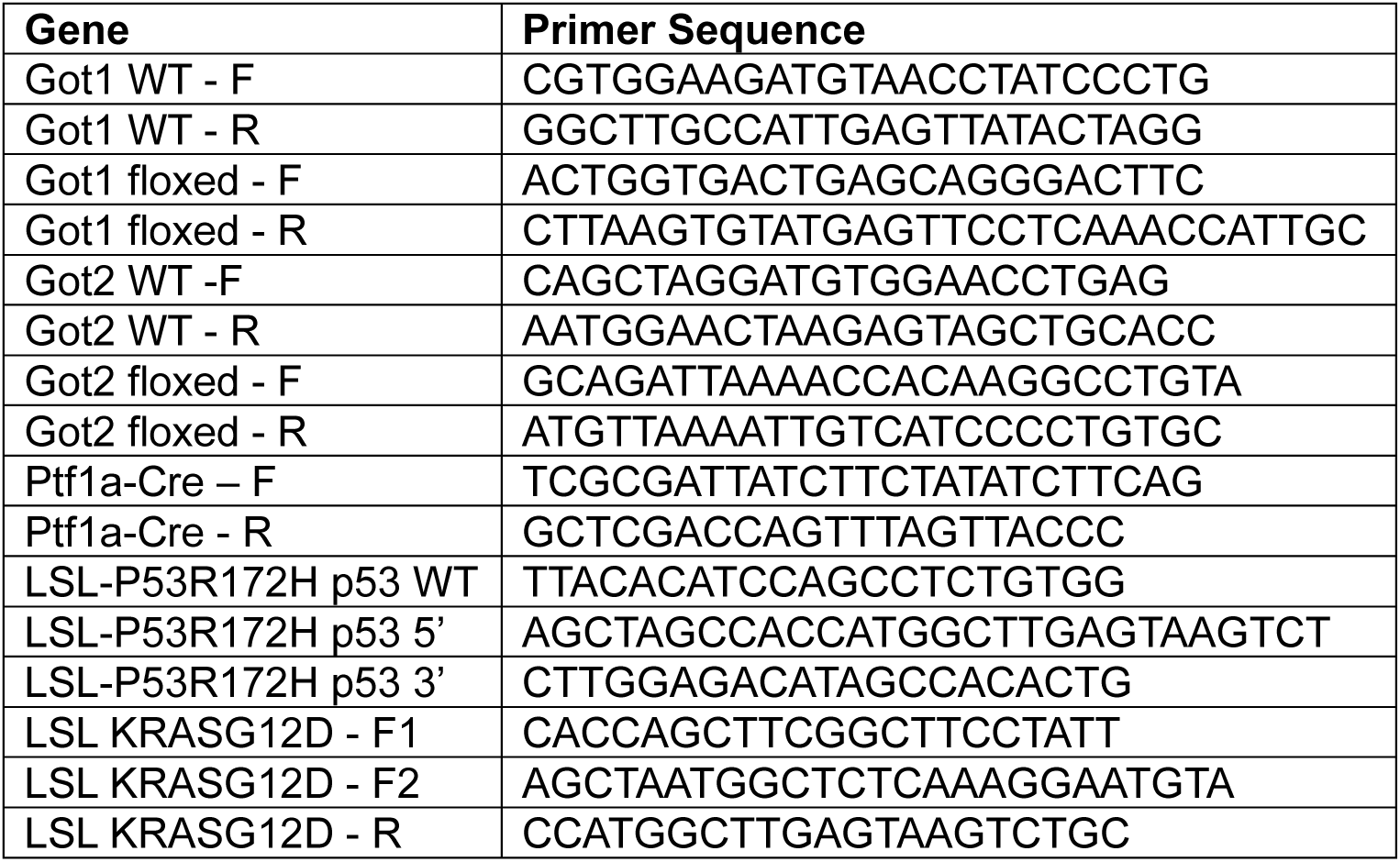
PCR primers for genotyping.

## Statistics and reproducibility

**Figure 1: Mouse pancreatic functions are maintained without Got1 or Got2.**

**C,** Statistical significance was measured using two-tailed t-tests. For pancreatic weight, p<0.0001. For body weight, p=0.0105. For pancreatic weight/body weight, p<0.0001. **E,** Statistical significance was measured using a two-tailed t-test. p<0.0001. **F,** Statistical significance was measured using a two-tailed t-test. p=0.7059. **G,** Statistical significance was measured using a two-tailed unpaired t-test. p=0.1455. **H,** Statistical significance was measured using a two-tailed unpaired t-test. p=0.0428. **I,** The area under the curve (AUC) was calculated for each individual mouse. Statistical significance was measured using a two-tailed unpaired t-test comparing the AUC of each group. p=0.6451. **J,** The area under the curve (AUC) was calculated for each individual mouse. Statistical significance was measured using a two-tailed unpaired t-test comparing the AUC of each group. p=0.1536. **K,** HOMA-IR was calculated by multiplying fasting glucose by fasting insulin and dividing by 22.5. Statistical significance was measured using a two-tailed unpaired t-test. p=0.1335. **L,** The glucose insulin index was calculated by multiplying the area under the curve of glucose and the area under the curve of insulin and 10^-6^. Statistical significance was measured using a two-tailed unpaired t-test. p=0.4504. **M,** Statistical significance was measured using two-tailed unpaired t-tests. No serum control (2 wells of reagent only); Cerulein-treated wildtype mice (2 wells per mouse, n=4 mice); *Got1^f/f^;Got2^f/f^*Cre negative mice (2 wells per mouse, n=5 mice); *Ptf1a-Cre*;*Got1^f/f^;Got2^f/f^*mice (2 wells per mouse, n=2 mice). Comparing cerulein-treated to DKO, p <0.000001****. Comparing cerulein-treated to Cre- mice, p<0.000001****. Comparing DKO to Cre- mice, p=0.000030****. **N-Q,** Statistical significance was measured using two-tailed unpaired t-tests. **N,** Aspartate, p=0.0464*. **O,** glutamate, p=0.6654. **P,** α-ketoglutarate, p=0.4040. **Q,** lactate/pyruvate, p=0.2906.

Figure 2: **Knockout of pancreatic Got1 or Got2 does not impact precancerous lesion formation.**

**B,** Statistical significance was measured with a two tailed unpaired t-test. Pancreas weight, p=0.8877. Body weight, p=0.8860. Pancreas/body weight, p=0.7337. **C,** Statistical significance was measured with a one-way ANOVA with Dunnet’s multiple comparisons. pancreas weight, KC to KC;*Got2^f/+^*, p=0.3368. KC to KC;*Got2^f/f^,* p=0.3735. Body weight, KC to KC;*Got2^f/+^*, p=0.0603. KC to KC;*Got2^f/f^,* p=0.0370*. Pancreas/body weight, KC to KC;*Got2^f/+^*, p=0.7527. KC to KC;*Got2^f/f^,* p=0.9520. **D,** Statistical significance was measured with a one-way ANOVA with Dunnet’s multiple comparisons. pancreas weight, KC to KC;*Got1^f/+^*, p=0.9569. KC to KC;*Got1^f/f^,* p=0.3786. Body weight, KC to KC;*Got1^f/+^*, p=0.3034. KC to KC;*Got1^f/f^,* p=0.6767. Pancreas/body weight, KC to KC;*Got1^f/+^*, p=0.9738. KC to KC;*Got1^f/f^,* p=0.5312. **E,** Statistical significance was measured with a one-way ANOVA with Dunnet’s multiple comparisons. pancreas weight, KC to KC;*Got2^f/+^*, p=0.9412. KC to KC;*Got2^f/f^,* p=0.8980. Body weight, KC to KC;*Got2^f/+^*, p=0.9432. KC to KC;*Got2^f/f^,* p=0. 9342. Pancreas/body weight, KC to KC;*Got2^f/+^*, p=0.9178. KC to KC;*Got2^f/f^,* p=0.9280. **F,** A KC;*Got2^f/f^* 12-month mouse was removed from the grading analysis after using Grubb’s outlier test with an alpha of 0.2. Statistical significance was measured using one-way ANOVAs. Acinar, p=0.3400. ADM, p=0.2583. PanIN 1A, p=0.2740. PanIN 2, p=0.5996. PanIN 3, p=0.7651. PDA, p=0.1406.

**Figure 3: Loss of Got1 or Got2 does not improve survival in genetically engineered models of pancreatic cancer.**

**B,** Statistical significance of survival time was measured using a one-way ANOVA, p= 0.4689. Average survival time: KPC (n=6) 134.3 days, KPC;*Got1^f/+^* het (n=11) 128.2 days, KPC;*Got1^f/f^* null (n=5) 145.8 days. **C,** Statistical significance of survival time was measured using a one-way ANOVA, p= 0.7353. Average survival time: KPC (n=6) 134.3 days, KPC;*Got2^f/+^* het (n=12) 133.5 days, KPC;*Got2^f/f^*143.7 days.

**Figure 4: Got1 or Got2 loss does not alter the composition of immune and epithelial compartments in the tumor microenvironment.**

**B,** Analysis performed on KPC, KPC;*Got1^f/+^*, and KPC;*Got1^f/f^* pancreata (n=5 mice per group). Statistical significance was measured with one-way ANOVAs with Dunnet’s multiple comparisons. CD3+/CD8-/FoxP3-; KPC to KPC;*Got1^f/+^*, p=0.4200. KPC to KPC;*Got1^f/f^*, p=0.8784. **C,** CD3+/CD8-/FoxP3+; KPC to KPC;*Got1^f/+^*, p=0.9916. KPC to KPC;*Got1^f/f^*, p=0.201*. **D,** CD3+/CD8+/FoxP3-; KPC to KPC;*Got1^f/+^*, p=0.7204. KPC to KPC;*Got1^f/f^*, p=0.4894. **E,** Total T Cells, KPC to KPC;*Got1^f/+^*, p=0.5094. KPC to KPC;*Got1^f/f^*, p=0.9351. **F,** F480+/PDL1-; KPC to KPC;*Got1^f/+^*, p=0.7382. KPC to KPC;*Got1^f/f^*, p=0.6963. **G,** F480+/PDL1+; KPC to KPC;*Got1^f/+^*, p=0.3500. KPC to KPC;*Got1^f/f^*, p=0.5652. **H,** F480+; KPC to KPC;*Got1^f/+^*, p=0.7376. KPC to KPC;*Got1^f/f^*, p=0.6958.**I,** CK19+/PDL1-; KPC to KPC;*Got1^f/+^*, p=0.9953. KPC to KPC;*Got1^f/f^*, p=0.5750. **J,** CK19+/PDL1+; KPC to KPC;*Got1^f/+^*, p=0.8662. KPC to KPC;*Got1^f/f^*, p=0.9946.**K,** CK19+; KPC to KPC;*Got1^f/+^*, p=0.9893. KPC to KPC;*Got1^f/f^*, p=0.5742. **L,** Analysis performed on KPC, KPC;*Got2^f/+^,* and KPC;*Got2^f/f^* pancreata (n=5 mice per group). Statistical significance was measured with one-way ANOVAs with Dunnet’s multiple comparisons. CD3+/CD8-/FoxP3-; KPC to KPC;*Got2^f/+^,* p=0.9672. KC to KPC;*Got2^f/f^*, p=0.5817. **M,** CD3+/CD8-/FoxP3+; KPC to KPC;*Got2^f/+^,* p=0.0084**. KC to KPC;*Got2^f/f^*, p= 0.8722. **N,** CD3+/CD8+/FoxP3-; KPC to KPC;*Got2^f/+^,* p=0.2592. KC to KPC;*Got2^f/f^*, p=0.8374. **O,** Total T Cells; KPC to KPC;*Got2^f/+^,* p=0.8970. KC to KPC;*Got2^f/f^*, p=0.6198.**P,** F480+/PDL1-; KPC to KPC;*Got2^f/+^,* p=0.5403. KC to KPC;*Got2^f/f^*, p=0.7421.**Q,** F480+/PDL1+; KPC to KPC;*Got2^f/+^,* p=0.9665. KC to KPC;*Got2^f/f^*, p=0.1716. **R,** F480+; KPC to KPC;*Got2^f/+^,* p=0.5404. KC to KPC;*Got2^f/f^*, p=0.7414. **S,** CK19+/PDL1-; KPC to KPC;*Got2^f/+^,* p=0.2352. KC to KPC;*Got2^f/f^*, p=0.3086. **T,** CK19+/PDL1+; KPC to KPC;*Got2^f/+^,* p=0.9479. KC to KPC;*Got2^f/f^*, p=0.7145. **U,** CK19+; KPC to KPC;*Got2^f/+^,* p=0.2715. KC to KPC;*Got2^f/f^*, p=0.3132

**Figure 5: PDA cells adapt their metabolism to counter GOT loss.**

**B,** Statistical significance was measured with a one-way ANOVA with Dunnett’s multiple comparisons. sgNT to Got1 sg1, p=0.9967. sgNT to Got2 sg1, p=0.5040. sgNT to Got2 sg2, p=0.5915. **C,** Statistical significance was measured with a one-way ANOVA with Dunnett’s multiple comparisons. sgNT to Got1 sg1, p=0.9951. sgNT to Got2 sg1, p=0.4177. **F-M,** Statistical significance was measured with a one-way ANOVA with Dunnet’s multiple comparisons. **F,** Aspartate; MT3: NT to Got1 sg1, 0.0030**. NT to Got2 sg1, 0.0847. **G,** 7940B: NT to Got1 sg1, p=0.0564. NT to Got2 sg1, p=0.0681. NT to Got2 sg2, p=0.1598. **H,** Glutamate; MT3: NT to Got1, 0.5402, NT to Got2 sg1, 0.0081**. **I,** 7940B: NT to Got1 sg1, 0.1594. NT to Got2 sg1, 0.5562. NT to Got2 sg2, 0.3036. **J,** αKG; MT3: NT to Got1 sg1, p= 0.4166. NT to Got2 sg1, p=0.2319. **K,** 7940B: NT to Got1 sg1, p=0.3497. NT to Got2 sg1, p=0.3497. NT to Got2 sg2, p=0.1365. **L,** Lactate/pyruvate; MT3: NT to Got1 sg1, p= 0.<0.0001****. NT to Got2 sg1, p=0.0382*. **M,** 7940B: NT to Got1 sg1, p=0.0113*. NT to Got2 sg1, p=0.0380*. NT to Got2 sg2, p=0.0309*. **N,** Statistical significance was measured with two tailed unpaired t-tests (n=5 mice per group). Pancreas weight, p= 0.8230. body weight, p= 0.4937. Pancreas/body weight, p= 0.6871. **O,** Statistical significance was measured with one-way ANOVAs (n=5 mice per group) with Dunnett’s multiple comparisons. Pancreas weight, sgNT to Got2 sg1, p=0.9904. sgNT to Got2 sg2, p=0.9954. Body weight, sgNT to Got2 sg1, p=0.4571. sgNT to Got2 sg2, p=0.6886. Pancreas/Body weight, sgNT to Got2 sg1, p=0.8335. sgNT to Got2 sg2, p=0.9609.

**Supplementary Figure 1**

**B,** Heatmap was generated identifying metabolites with a p-value<0.1 compared to the control sample (Cre negative mice). Samples included pancreata of 8-week old *Got1^f/f^;Got2^f/f^* (n=4) or *Ptf1a-Cre*;*Got1^f/f^;Got2^f/f^*(n=3) mice

**Supplementary Figure 2**

**A,** Statistical significance was measured with a one-way ANOVA with Dunnet’s multiple comparisons. Pancreas weight, KPC to KPC;*Got1^f/f^,* p=0.6485. KPC to KPC;*Got1^f/f^*, p=0.3787. Body weight, KPC to KPC;*Got1^f/f^,* p=0.9860. KPC to KPC;*Got1^f/f^*, p=0.3603. Pancreas/Body weight, KPC to KPC;*Got1^f/f^,* p=0/7897. KPC to KPC;*Got1^f/f^*, p=0.6374. **B,** Statistical significance was measured with a one-way ANOVA with Dunnet’s multiple comparisons. Pancreas weight, KPC to KPC;*Got2^f/+^,* p=0.7890. KPC to KPC;*Got2^f/f^*, p=0.9039. Body weight, KPC to KPC;*Got2^f/+^,* p=0.9974. KPC to KPC;*Got2^f/f^*, p=0.3601. Pancreas/Body weight, KPC to KPC;*Got2^f/+^,* p=0.7340. KPC to KPC;*Got2^f/f^*, p=0.9923.

**Supplementary Figure 3**

Heatmap was generated identifying metabolites with a p-value<0.1 compared to the control sample (sgNT cell lines). n=3 biologically independent samples per group.

**Supplementary Figure 4**

Heatmap was generated identifying metabolites with a p-value<0.1 compared to the control sample (sgNT cell lines). n=3 biologically independent samples per group.

**Supplementary Figure 5**

Heatmaps were generated identifying metabolites with a p-value<0.1 compared to the control sample (sgNT cell lines). n=3 biologically independent samples per group, n=2 for Got2 sg2.

**Supplementary Figure 6**

Heatmaps were generated identifying metabolites with a p-value<0.1 compared to the control sample (sgNT cell lines). n=3 biologically independent samples per group, n=2 for Got2 sg2.

**Supplementary Figure 1.**
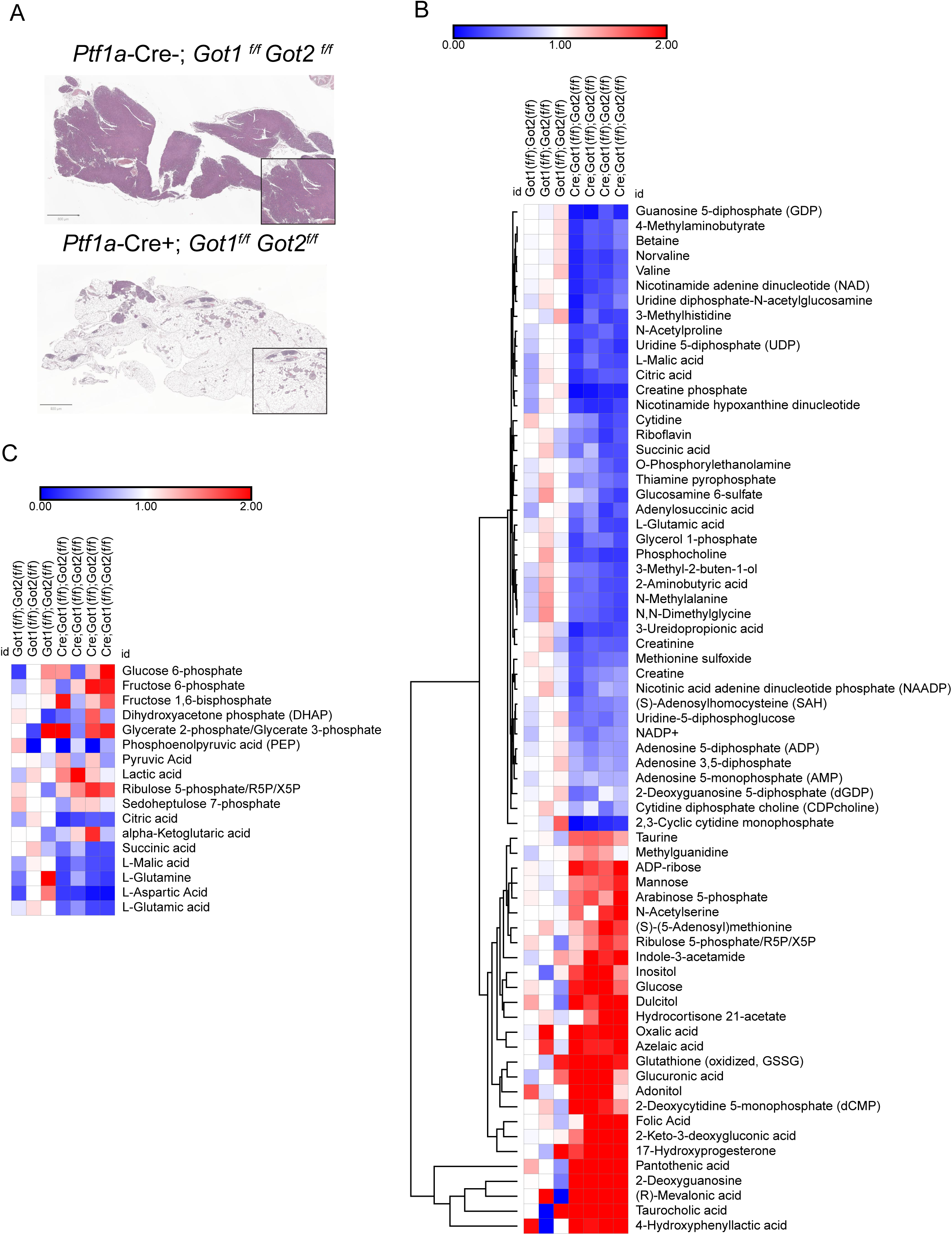
**A,** Representative hematoxylin and eosin (H&E) staining of *Got1^f/f^;Got2^f/f^*(Cre negative) and *Ptf1a-Cre*;*Got1^f/f^;Got2^f/f^*pancreata. Scale bar is 800µm. **B,** Unsupervised hierarchical clustering of differentially abundant metabolites (p<0.1, compared to Cre negative mice) in pancreata of 8-week old *Got1^f/f^;Got2^f/f^* (n=3) or *Ptf1a-Cre*;*Got1^f/f^;Got2^f/f^* (n=4) mice, as measured by LC-MS/MS. **C,** Curated list of glycolytic, pentose phosphate, and TCA cycle intermediate metabolites from the dataset generated in **B**.

**Supplementary Figure 2.**
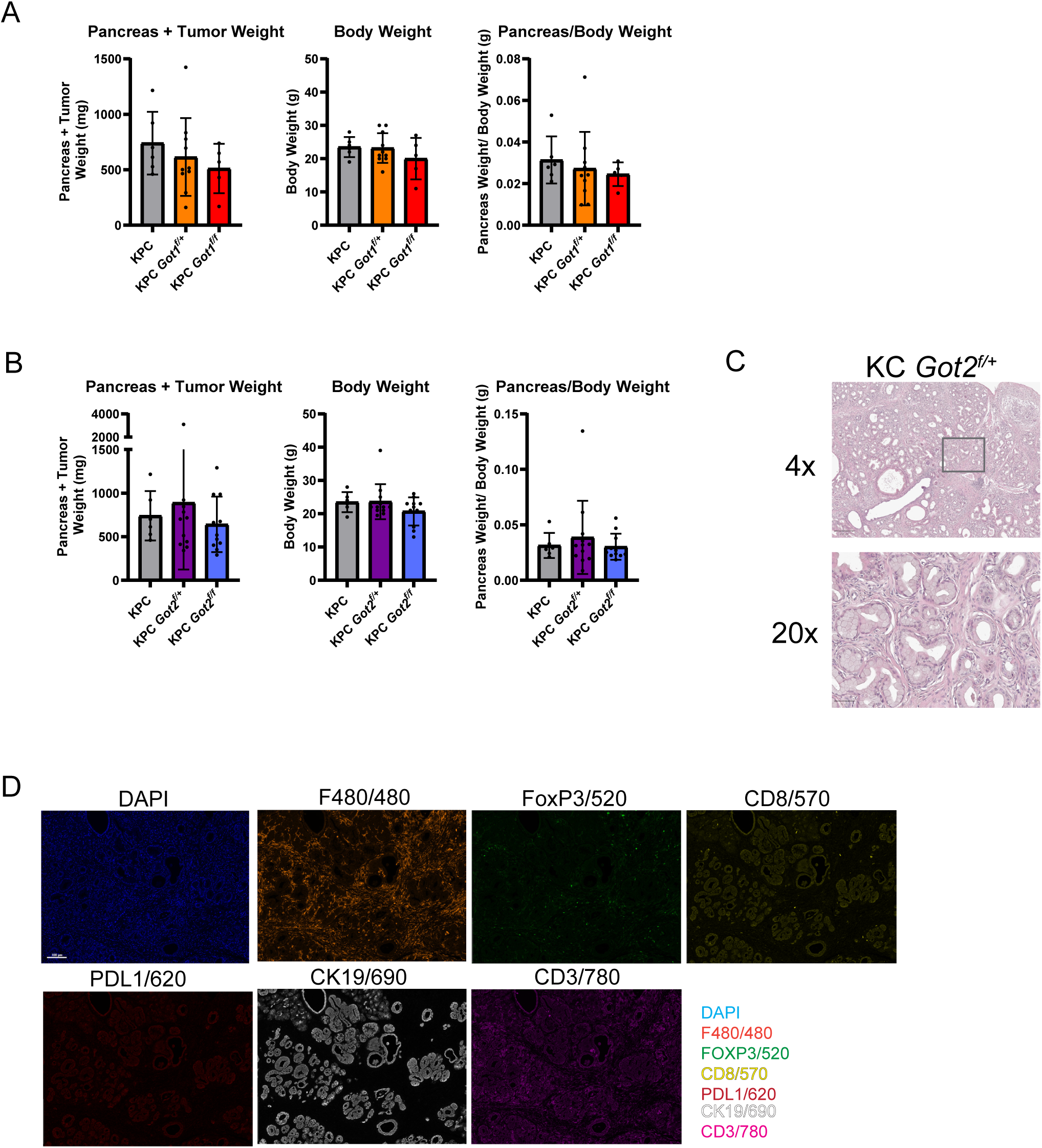
**A,** Tumor and body weight of KPC (n=6), KPC;*Got1^f/+^* (n=11), and KPC;*Got1^f/f^* (n=5) mice. **B,** Tumor and body weight of KPC (n=6), KPC;*Got2^f/+^* (n=12), and KPC;*Got2^f/f^* (n=11) mice. **C,** Representative hematoxylin and eosin (H&E) staining of 12-month-aged KC; *Got2^f/+^* pancreata. Scale bar is 100µm. **D,** Individual immunofluorescence panels for multiplex IF. Scale bar is 250µm for 4x and 50µm for 20x images.

**Supplementary Figure 3.**
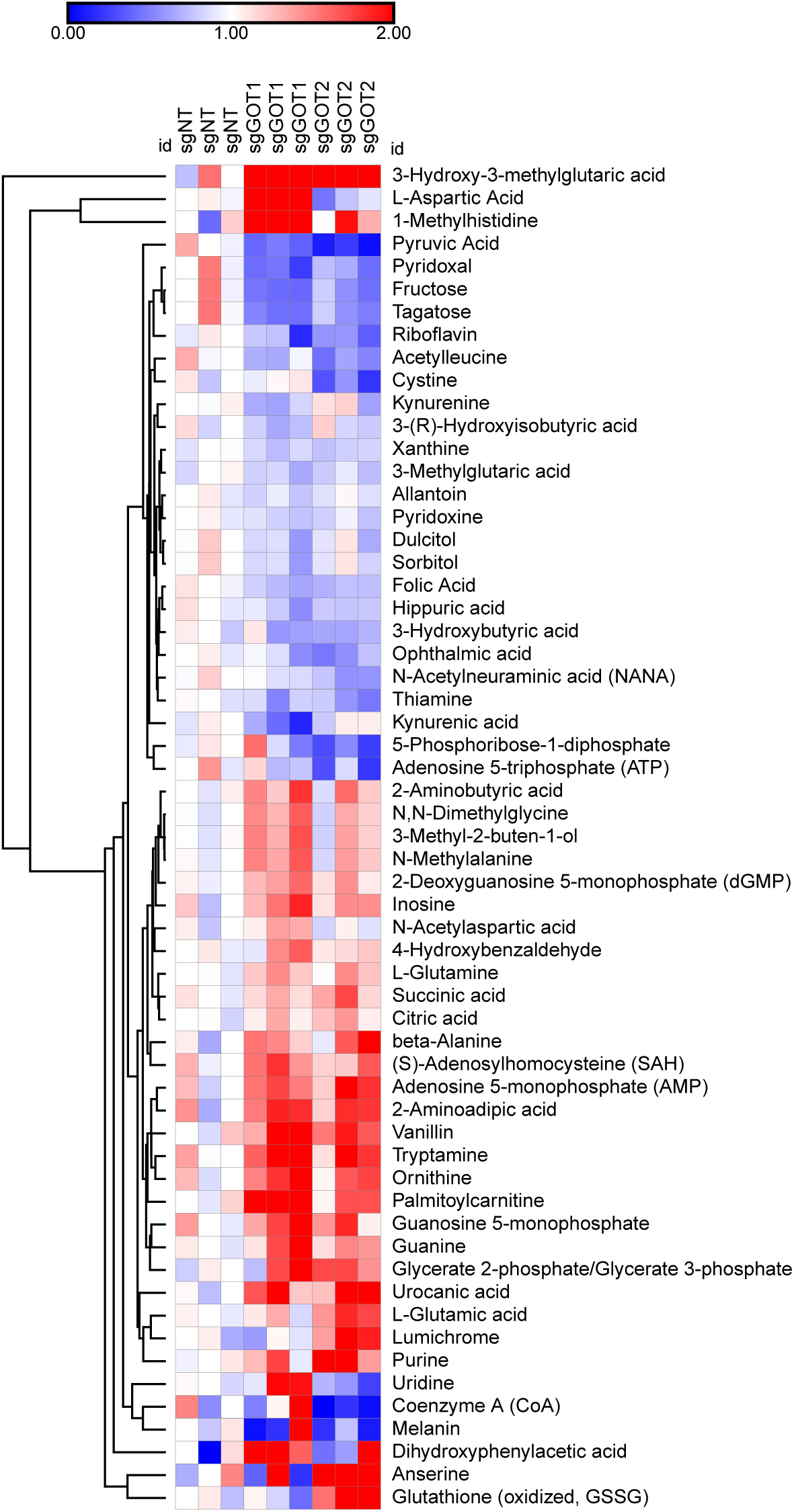
Unsupervised hierarchical clustering of differentially abundant intracellular metabolites (p<0.1, compared to sgNT) from MT3 cells with *Got1* or *Got2* knockout as measured by LC-MS/MS. n=3 biologically independent samples per group.

**Supplementary Figure 4.**
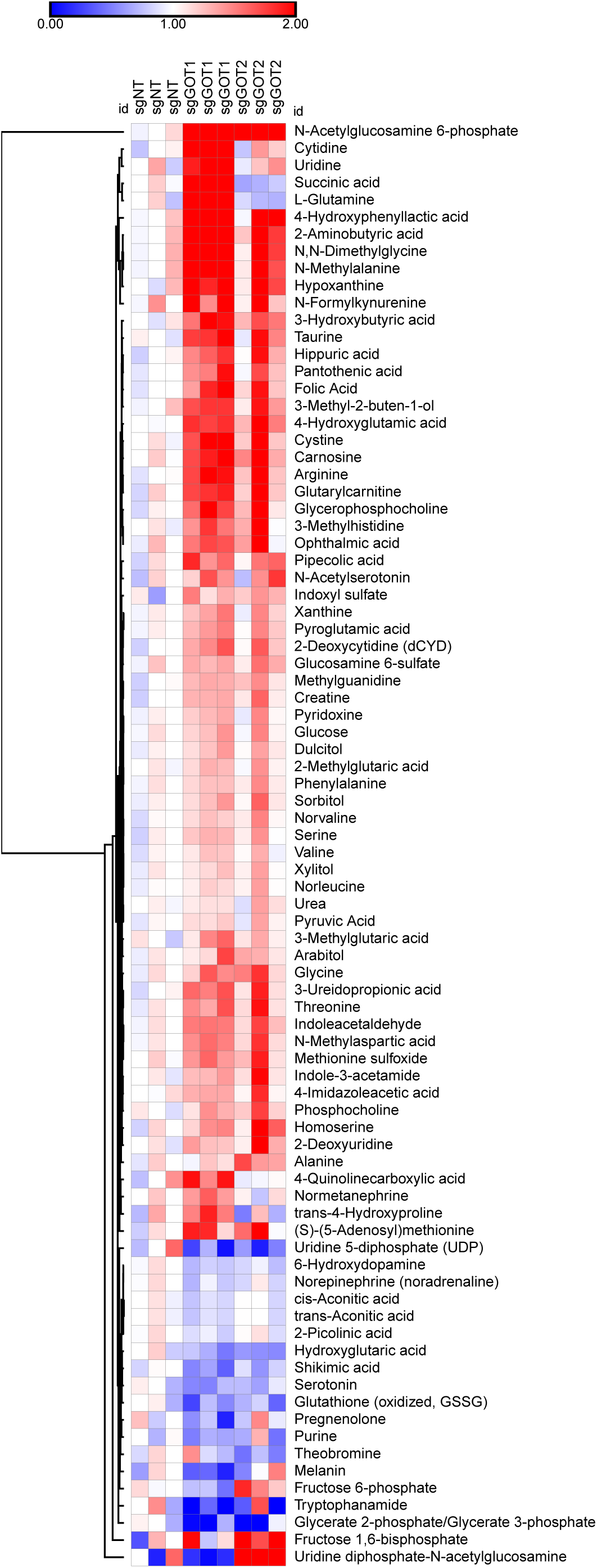
Unsupervised hierarchical clustering of differentially abundant extracellular metabolites (p<0.1, compared to sgNT) from MT3 cells with *Got1* or *Got2* knockout as measured by LC-MS/MS. n=3 biologically independent samples per group.

**Supplementary Figure 5.**
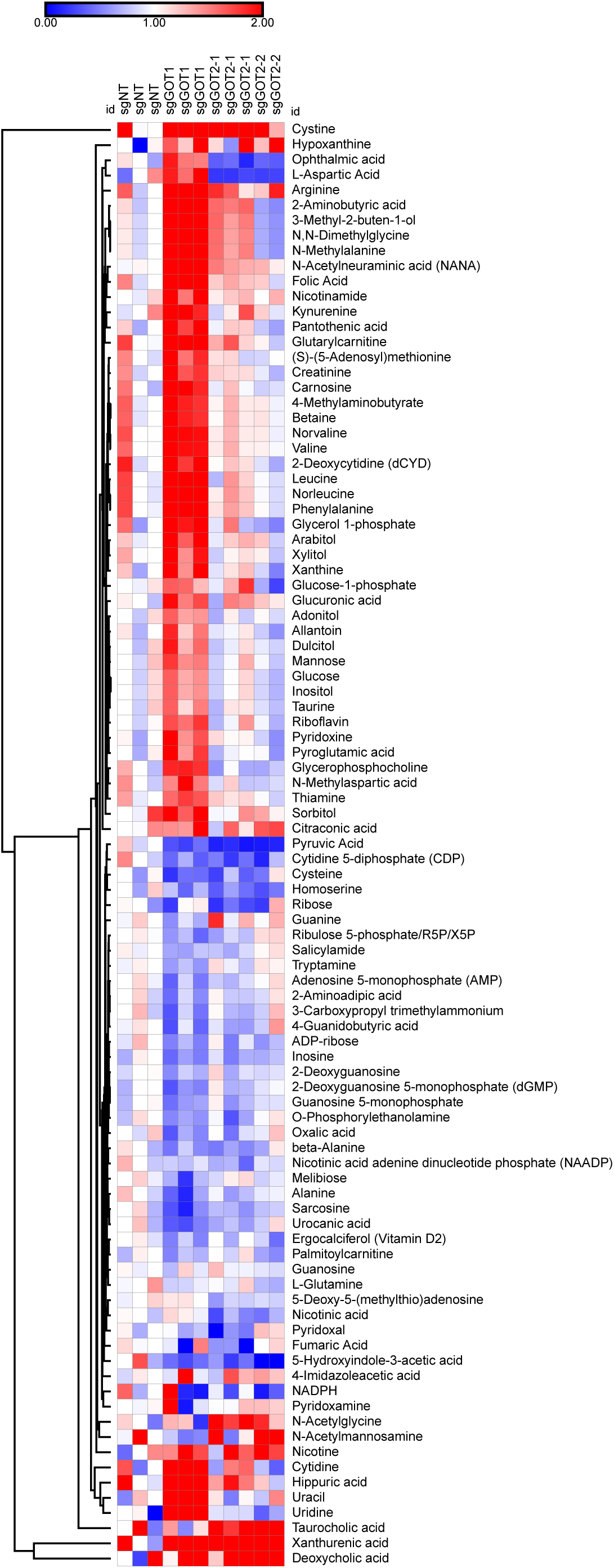
Unsupervised hierarchical clustering of differentially abundant intracellular metabolites (p<0.1, compared to sgNT) from 7940B cells with *Got1* or *Got2* knockout as measured by LC-MS/MS. n=3 biologically independent samples per group; n=2 for Got2 sg2.

**Supplementary Figure 6.**
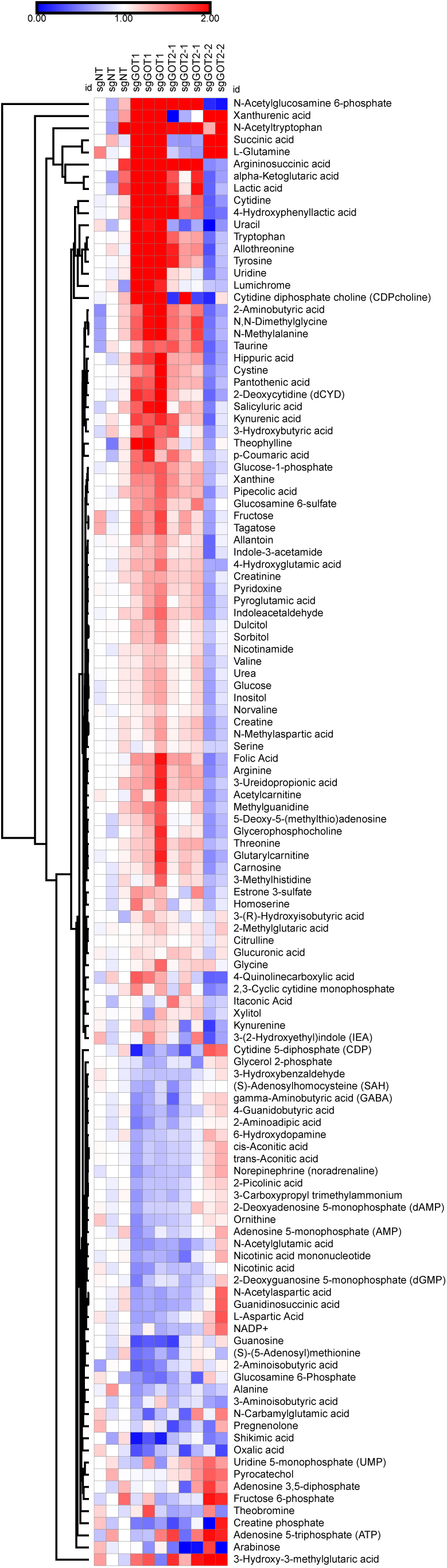
Unsupervised hierarchical clustering of differentially abundant extracellular metabolites (p<0.1, compared to sgNT) from 7940B cells with *Got1* or *Got2* knockout as measured by LC-MS/MS. n=3 biologically independent samples per group; n=2 for Got2 sg2.

